# Molecular Determinants of μ-Conotoxin KIIIA Interaction with the Human Voltage-Gated Sodium Channel NaV1.7

**DOI:** 10.1101/654889

**Authors:** Ian H. Kimball, Phuong T. Nguyen, Baldomero M. Olivera, Jon T. Sack, Vladimir Yarov-Yarovoy

**Affiliations:** Department of Physiology and Membrane Biology, UC Davis, Davis, CA, USA; Department of Anesthesiology and Pain Medicine, UC Davis, Davis, CA, USA; Department of Biology, University of Utah, Salt Lake City, UT, USA

## Abstract

The voltage-gated sodium (Na_V_) channel subtype Na_V_1.7 plays a critical role in pain signaling, making it an important drug target. Here we studied the molecular interactions between μ-conotoxin KIIIA (KIIIA) and the human Na_V_1.7 channel (hNa_V_1.7). We developed a structural model of hNa_V_1.7 using Rosetta computational modeling and performed in silico docking of KIIIA using RosettaDock to predict residues forming specific pairwise contacts between KIIIA and hNa_V_1.7. We experimentally validated these contacts using mutant cycle analysis. Comparison between our KIIIA-hNa_V_1.7 model and the cryo-EM structure of KIIIA-hNa_V_1.2 revealed key similarities and differences between Na_V_ channel subtypes with potential implications for the molecular mechanism of toxin block. The accuracy of our integrative approach, combining structural data with computational modeling, experimental validation, and molecular dynamics simulations, suggests that Rosetta structural predictions will be useful for rational design of novel biologics targeting specific Na_V_ channels.

## Introduction

Voltage-gated sodium (Na_V_) channels play a key role in the action potential generation in excitable cells (Hille, 2001;Catterall, 2014;Ahern et al., 2016). The nine subtypes of Na_V_ channel α-subunits (named Na_V_1.1-Na_V_1.9) are differentially expressed throughout tissues, and are targets of therapeutics for pain, cardiac arrhythmias, and epilepsy (Catterall et al., 2005). Human Na_V_1.7 (hNa_V_1.7) channel is important for pain signaling and its mutations have been linked to severe pain disorders ranging from complete lack of pain sensation to extreme sensitivity to pain (Dib-Hajj et al., 2013;Bennett et al., 2019;Dib-Hajj and Waxman, 2019). Clinical use of local anesthetic drugs, such as lidocaine, is limited because they bind to a highly conserved receptor site within the Na_V_ channel pore lumen, and are consequently non-selective among human Na_V_ subtypes (Ragsdale et al., 1994;Yarov-Yarovoy et al., 2001;Yarov-Yarovoy et al., 2002;Nguyen et al., 2019). The receptor sites of other Na_V_ blockers have nuanced differences between subtypes and molecular mechanisms of channel blockade that could enable rational design of Na_V_ subtype-selective therapeutics (Payandeh and Hackos, 2018).

The search for novel Na_V_ channel modulators has identified small disulfide-knotted peptide toxins from cone snails (conotoxins) (Wilson et al., 2011), which target the extracellular vestibule of the Na_V_ channel pore and offer useful peptide scaffolds for rational design of novel peptide-based therapeutics to potentially treat pain, arrhythmias, and epilepsy (French et al., 2010;Gilchrist et al., 2014). μ-conotoxin KIIIA (KIIIA) is a 16 amino acid peptide that potently inhibits TTX-sensitive Na_V_ channels (Zhang et al., 2007;Wilson et al., 2011;Khoo et al., 2012) (Figure 1A). A pain assay in mice suggested that KIIIA potentially has analgesic properties (Zhang et al., 2007). Notably, a variant of saxitoxin that targets the same receptor site in hNav1.7 as KIIIA is now in a Phase I clinical trial for treatment of post-operative pain (Mulcahy et al., 2019;Pajouhesh et al., 2020;SiteOne Therapeutics, 2021). KIIIA has variable degrees of affinity and block for the different Na_V_ channel subtypes, with 5 nM affinity for rat Na_V_1.2, 37 nM for rat Na_V_1.4, and 97 nM for hNa_V_1.7 (Zhang et al., 2007;McArthur et al., 2011;Wilson et al., 2011). Structure-activity relationship studies have identified the KIIIA residues K7, W8, R10, D11, H12 and R14 as key for binding to various Na_V_ channel subtypes (Zhang et al., 2007;McArthur et al., 2011). Specifically, K7, R10 and R14 have been shown to contribute to both binding affinity and block of hNa_V_1.7 (McArthur et al., 2011). Notably, the relative contribution of KIIIA residues in binding to Na_V_ channels vary between channel subtypes. For example, substitution R14A in KIIIA reduces the affinity for Na_V_1.2 and Na_V_1.4 by 740-fold and 180-fold, respectively, while reducing the affinity for Na_V_1.7 by only 5-fold (McArthur et al., 2011). Similarly, substitution R10A in KIIIA reduces the affinity for Na_V_1.2 and Na_V_1.4 by 32-fold and 27-fold, respectively, while reducing the affinity for Na_V_1.7 by only 14-fold (McArthur et al., 2011). In addition, KIIIA blocks Na_V_ channels incompletely and can co-bind with tetrodotoxin (TTX) to TTX-sensitive Na_V_ channels (Zhang et al., 2009).

**Figure 1.**
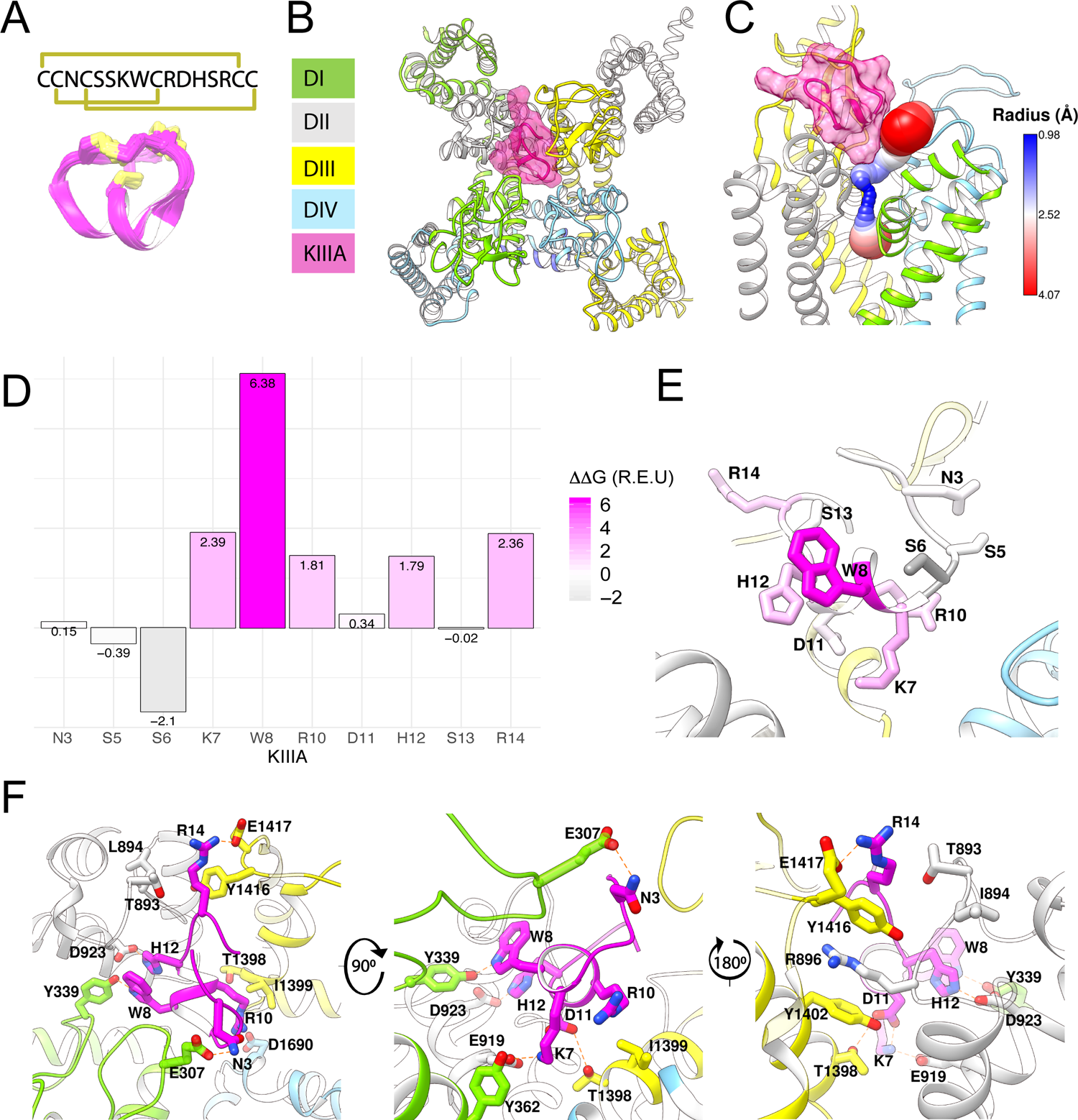
Structural model of KIIIA - hNa_V_1.7 features eccentric binding of toxin to the outer pore. (**A**) Structure and sequence of KIIIA (PDB: 2LXG) (Khoo et al., 2012) show an alpha helical core stabilized by three disulfide bridges. (**B**) Extracellular view of our KIIIA – hNaV1.7 complex homology model based on the EeNa_V_1.4-β1 cryo-EM structure (Yan et al., 2017). Channel domains are depicted according to color keys, and KIIIA is shown in magenta ribbon and surface. KIIIA binds eccentrically to the outer pore between DII and DIII. (**C**) A side view of channel pore reveals an incomplete block of KIIIA with KIIIA bound (magenta) and cavity volume. (**D**) Rosetta alanine scan identified residues K7, W8, R10, H12, and R14 as significant contributors to binding energy. (**E**) Heatmap of Rosetta ΔΔG on KIIIA structure shows the importance of the helical region for binding. (**F**) Close-up views of key interactions at KIIIA – hNa_V_1.7 interface.

A previous study identified the importance of Na_V_ channel residues near the selectivity filter on the P2-helix in domain III (DIII) for their apparent coupling to residues R10 and R14 on KIIIA (McArthur et al., 2011). Notably, the P2-helix in DIII of hNa_V_1.7 has threonine at position 1398 and an isoleucine at position 1399, while all other human Na_V_ channels have methionine and aspartate at the corresponding positions (McArthur et al., 2011). These residues were proposed to play an important role in the selectivity of KIIIA binding to Na_V_1.2 and Na_V_1.4 versus to Na_V_1.7 (McArthur et al., 2011). Molecular modeling of KIIIA binding to rNa_V_1.4 using restraints from experimental data suggested specific contacts between KIIIA and the P2-helix in DIII (Korkosh et al., 2014). However, these studies did not provide an explanation for the significant effect of the KIIIA mutations H12A, W8A and D11A on toxin affinity.

In this study, we used computational and experimental approaches to investigate the molecular mechanism of the KIIIA interaction with hNa_V_1.7. We specifically focused on human Na_V_1.7 as a target due to its importance in pain signaling and our goal to rationally design novel KIIIA-based peptides targeting human Na_V_1.7 as potential therapeutics to treat chronic pain. We selected KIIIA among conotoxins identified to date because it has the highest potency for human Na_V_1.7 (Zhang et al., 2007;McArthur et al., 2011). We present a structural model of KIIIA binding to the hNa_V_1.7 channel based on the eukaryotic electric eel Na_V_1.4 cryo-EM structure (Yan et al., 2017). Our model revealed binding of KIIIA to hNa_V_1.7 at the interface between the P2-helices in domain II (DII) and DIII, which exposed a partially open ion conduction pathway that may explain the incomplete blocking characteristic of the toxin. While many KIIIA mutations have been previously characterized on Nav1.2 and Nav1.4 channels, only a limited number of KIIIA mutants had been tested on Nav1.7. We independently characterized many of the KIIIA mutations on Nav1.7 in our laboratory, to confirm previously published data and to validate affinities that serve as the basis of mutant cycle calculations. Our study for the first time tested effects of the following mutations: (1) KIIIA W8 and D11A on the wild-type hNav1.7; (2) hNav1.7 Y362C, E919Q, D923A with the wild-type KIIIA; (3) paired toxin—channel mutations K7A—E919Q, D11A—E919Q, and H12A—D923A. We identified several unique contacts between KIIIA and extracellular loops on hNa_V_1.7, providing key structural insights into binding specificity for different Na_V_ channel subtypes. We used mutant cycle analysis to validate representative pairwise contacts between specific residues on KIIIA and hNa_V_1.7 identified from our structural model of the KIIIA – hNa_V_1.7 complex. Remarkably, the published cryo-EM structure of KIIIA - hNa_V_1.2 complex (Pan et al., 2019) agrees with findings from our computational modeling and functional study. The accuracy of peptide toxin – Na_V_ channel interaction modeling suggests that Rosetta predictions are sufficiently precise for the rational design of novel selective peptide inhibitors targeting Na_V_ channels with high selectivity and potency.

## Results

### Molecular modeling suggests eccentric binding of KIIIA to DII and DIII of the hNa_V_1.7 pore

To characterize the molecular mechanism of the KIIIA interaction with hNa_V_1.7, we utilized computational modeling and functional testing approaches as described below. When this study was conducted, the cryo-EM structure of the electric eel Na_V_1.4 (eeNa_V_1.4) (Yan et al., 2017) channel was the closest structural homolog available to build a homology model of hNa_V_1.7. The eeNa_V_1.4 structure shares ∼54% sequence identity with hNa_V_1.7 overall and ∼75% sequence identity over the hNa_V_1.7 pore region. We used the RosettaCM modeling approach (Song et al., 2013;Bender et al., 2016) to generate a structural model of hNa_V_1.7 based on the eeNa_V_1.4 structure (Yan et al., 2017) and the Rosetta protein-protein docking approach (Gray et al., 2003;Wang et al., 2007;Bender et al., 2016) to predict a structure of the KIIIA – hNa_V_1.7 complex and identify specific residues forming interactions between KIIIA and hNa_V_1.7 (see Materials and methods, and coordinates of our KIIIA – hNa_V_1.7 model in Supplement File – Model 1). Our model revealed an eccentric or off-center binding of KIIIA to hNa_V_1.7, where the KIIIA helical region is positioned off the central axis of the selectivity filter with the positively charged KIIIA residues facing the P2-helices (Figure 1B). The position and orientation of KIIIA in our model is different from KIIIA binding to the P2-helix in DIII previously suggested by computational modeling (McArthur et al., 2011;Korkosh et al., 2014) and lanthanide-based resonance energy transfer (Kubota et al., 2017) studies. The KIIIA binding site just above the selectivity filter in our model is different from TTX and saxitoxin (STX) binding deeper into the selectivity filter region (Shen et al., 2018;Shen et al., 2019). Mapping of the open aqueous space surrounding the KIIIA - hNa_V_1.7 binding interface revealed a tunnel traversing from the extracellular environment to the channel pore cavity (Figure 1C). The most constricted part of this aqueous tunnel is within the selectivity filter region, where the radius of the open aqueous space narrows to ∼1 Å. KIIIA bound to the upper region of the selectivity filter and constricted the open space to a minimum radius of ∼2.5 Å, which is large enough to allow sodium ion conduction and consistent with the characteristic incomplete block of Na_V_ channels by KIIIA (Zhang et al., 2007;McArthur et al., 2011). Notably, the cryoEM structure of KIIIA – hNa_V_1.2 (Pan et al., 2019) reveals a binding pose similar to our model, as described later in this paper (Figure 4A).

### Pairwise interactions identified from the KIIIA - hNa_V_1.7 complex model

To identify key KIIIA residues at the toxin – channel interface, we first examined the contribution of KIIIA residues to the interaction with hNa_V_1.7 using an *in silico* alanine scan. Non-cysteine residues on KIIIA were mutated to alanine and changes in Rosetta binding energy (ΔΔG) were reported in the arbitrary Rosetta Energy Units (R.E.U). Our analysis revealed the active surface of KIIIA with residues K7, W8, R10, H12, and R14 each having significant contribution to the binding energy (Figure 1D and E). K7, W8, R10, and H12 are located on the same face of KIIIA’s alpha helix, while R14 is located within the C-terminal tail region of the toxin. Our KIIIA – hNa_V_1.7 model predicts that positively charged residue K7 forms a salt bridge with E919 on the P2-helix in DII (we use hNa_V_1.7 residue numbering throughout the manuscript unless otherwise noted) (Figure 1F). In addition, W8 and H12 were shown to form hydrogen bonds with Y339 on the extracellular loop between S5 and the P1-helix (S5P1) in DI and D923 on the P2-helix in DII, respectively (Figure 1F). D11 is positioned near the interface between the P2-helices in DII and DIII and forms hydrogen bonds with both K7 on KIIIA and T1398 on the P2-helix in DIII (Figure 1F). The other positively charged KIIIA residues, R10 and R14, interact with two negatively charged residues: D1662 on the extracellular loop S5P1 in DIV and E1417 on the extracellular loop between the P2-helix and S6 (P2S6) in DIII. Notably, R14 also interacts with Y1416 on the extracellular P2S6 loop in DIII and contributes to a cation-π interaction tower formed by Y1402 on the P2-helix in DIII, R896 on the extracellular loop S5P1 in DII, and Y1416 (Figure 1F). The R10 of KIIIA is in proximity to I1399 on the P2-helix in DIII in agreement with the significant coupling energy between R10 and D1241 on the P2-helix in DIII in rNa_V_1.4 reported previously (McArthur et al., 2011). While KIIIA N3 is near E307 in the extracellular loop S5P1 in DI, this interaction may not be substantial as shown by minimal change in Rosetta binding energy (ΔΔG) from our *in silico* alanine scan (Figure 1D). N3 has been shown to be not critical for KIIIA interaction with rNa_V_1.2 and rNa_V_1.4 channels (Zhang et al., 2007).

### Functional mapping of KIIIA residues at the toxin – channel interface supports the predicted KIIIA – hNa_V_1.7 model

To test the accuracy of our KIIIA – hNa_V_1.7 model, we first confirmed the activity of the wild-type KIIIA on the wild-type hNa_V_1.7 using whole-cell voltage-clamp recordings. To estimate the KIIIA binding affinity, we performed concentration-response experiments and obtained an IC_50_ of 410±160 nM when fit to a Hill equation assuming a single binding site (Figure 2). This fitting suggested a maximal block of 95±3.3% of total current in agreement with previous studies concluding that Na_V_ channels retain 5-10% of their conductance when blocked by KIIIA (Zhang et al., 2007;McArthur et al., 2011). The WT-KIIIA first order association rate (*k_on_*) was determined based on single exponential fits to the kinetics of block after toxin addition (Table 1, see Materials and Methods equation 3). The extremely slow dissociation of the wild-type KIIIA from hNa_V_1.7 complicated accurate determination of dissociation kinetics, as less than 10% recovery was observed during wash-off experiments lasting up to ∼30 min. Constraining single exponential fits of the dissociation data to assume current recovers to initial levels, we obtained *k_off_* of 0.003 min^-1^ and a *K_d_* of 59 nM, which is close to the previously reported *K_d_* of 97 nM (McArthur et al., 2011). Temperature differences between our experiments (∼21°C) and the prior study (∼25°C) might be responsible for differences in kinetics and affinities.

**Figure 2.**
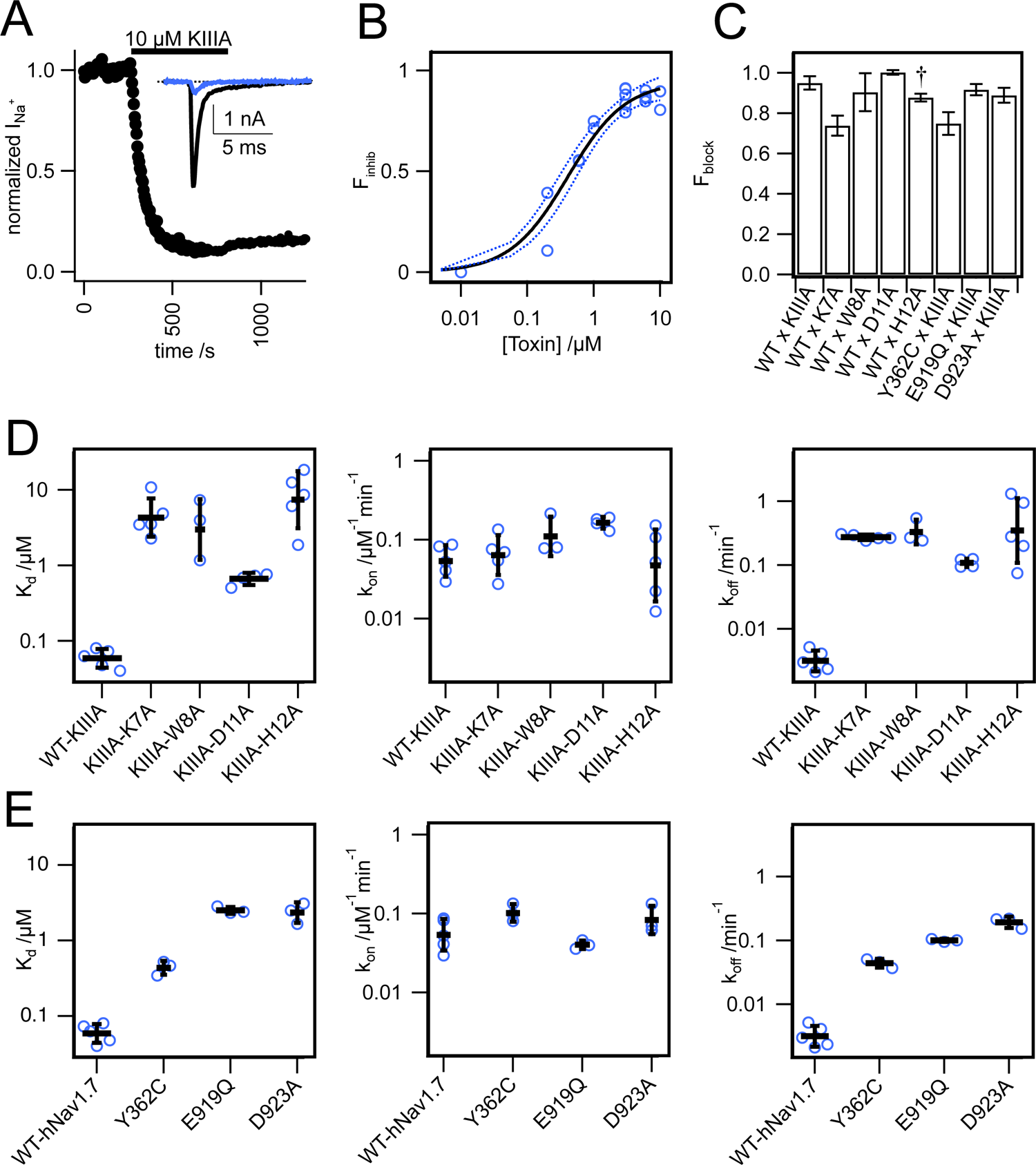
Functional studies of toxin variants and channel mutations. (**A**) Normalized peak I_Na_ from a whole cell voltage clamp experiment with 10 μM WT-KIIIA against WT-hNa_V_1.7, resulting in incomplete block and (inset) raw current traces before toxin (black) and after toxin (blue). (**B**) Hill-fit (black) and 95% confidence interval (dashed blue) of concentration-response data for WT-KIIIA against hNa_V_1.7 in HEK293 cells (IC_50_=0.41±0.16 μM mean±SD, n=2-4 cells per concentration), from maximum block recorded during association experiments. Empty circles represent single cells. **(C**) Calculated Fractional block (see Materials and methods) for toxin variants and channel mutants (mean±SEM). † WT x H12A block data were reported previously (McArthur et al., 2011). **(D**) Kinetic data from electrophysiological measurements show general agreement with Rosetta predicted energies. Alanine variants of residues K7, W8, D11, and H12 showed significant reductions in affinity (K_d_)(left), little change in association (*k_on_*)(middle), but marked increases in toxin dissociation (*k_off_*)(right). Bars are geometric mean±SD from n=3-5 cells per variant (reported in Table 1), and empty circles represent single cells. (**E**) Mutations to channel residues demonstrate reductions in affinity of the WT-KIIIA from Y362C, E919Q, and D923A (left), little change to toxin association (middle), and increases in dissociation (right), similar to the effects of toxin variants. Bars are geometric mean±SD from n=3-5 cells per variant (reported in table 2), and empty circles represent single cells.

**Table 1.**
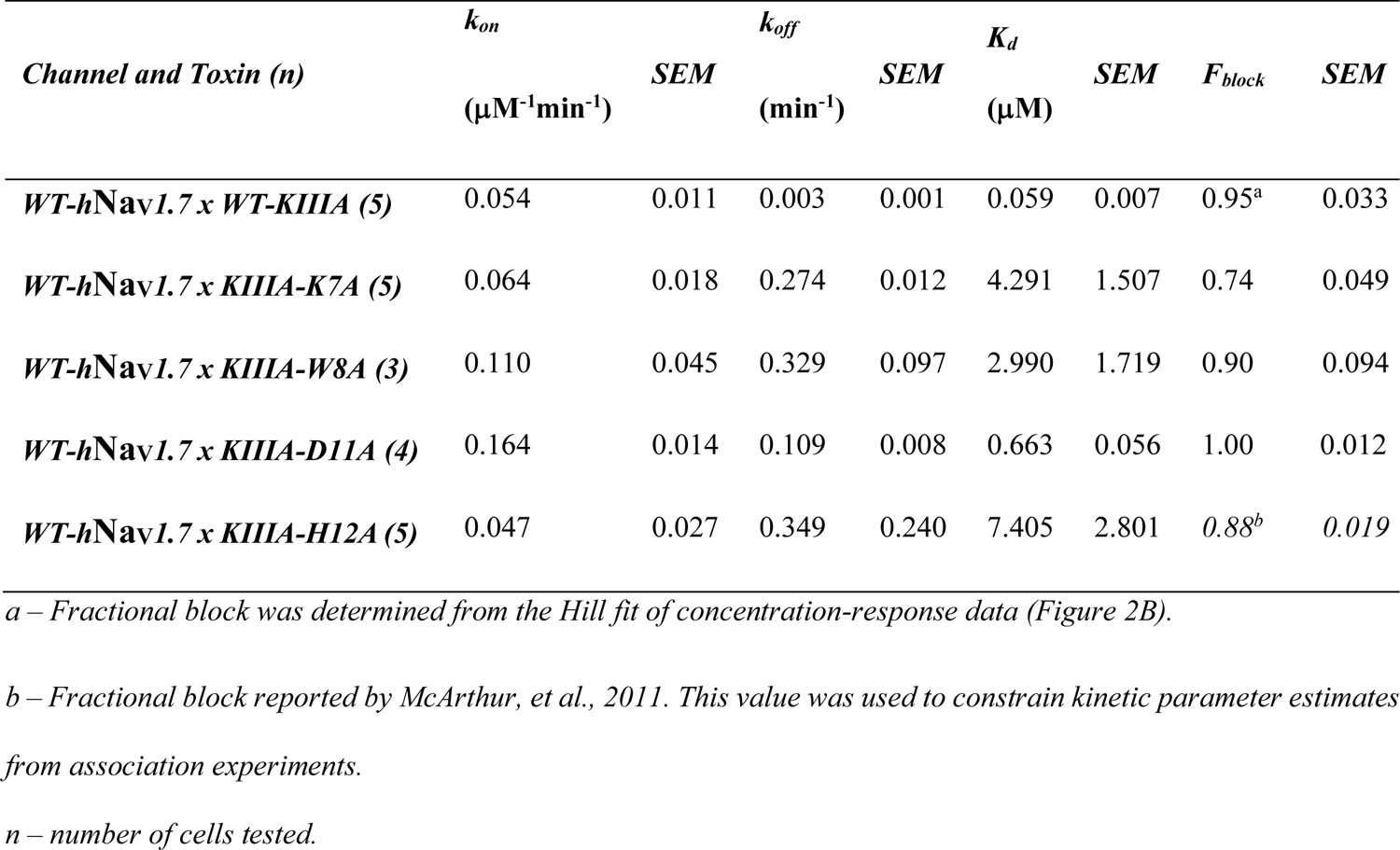
Block of hNa_V_1.7 by KIII variants

| *Channel and Toxin (n)* | $k_{on}$<br>($\mu\text{M}^{-1}\text{min}^{-1}$) | *SEM* | $k_{off}$<br>($\text{min}^{-1}$) | *SEM* | $K_d$<br>($\mu\text{M}$) | *SEM* | $F_{block}$ | *SEM* |
| --- | --- | --- | --- | --- | --- | --- | --- | --- |
| **WT-hNav1.7 x WT-KIIIA (5)** | 0.054 | 0.011 | 0.003 | 0.001 | 0.059 | 0.007 | 0.95<sup>a</sup> | 0.033 |
| **WT-hNav1.7 x KIIIA-K7A (5)** | 0.064 | 0.018 | 0.274 | 0.012 | 4.291 | 1.507 | 0.74 | 0.049 |
| **WT-hNav1.7 x KIIIA-W8A (3)** | 0.110 | 0.045 | 0.329 | 0.097 | 2.990 | 1.719 | 0.90 | 0.094 |
| **WT-hNav1.7 x KIIIA-D11A (4)** | 0.164 | 0.014 | 0.109 | 0.008 | 0.663 | 0.056 | 1.00 | 0.012 |
| **WT-hNav1.7 x KIIIA-H12A (5)** | 0.047 | 0.027 | 0.349 | 0.240 | 7.405 | 2.801 | 0.88<sup>b</sup> | 0.019 |

*a* – Fractional block was determined from the Hill fit of concentration-response data (Figure 2B).
*b* – Fractional block reported by McArthur, et al., 2011. This value was used to constrain kinetic parameter estimates from association experiments.
*n* – number of cells tested.

**Table 2.** Block of hNa_V_1.7 mutants by KIIIA

| *Channel and Toxin (n)* | *k<sub>on</sub>*<br>(μM<sup>-1</sup>min<sup>-1</sup>) | *SEM* | *k<sub>off</sub>*<br>(min<sup>-1</sup>) | *SEM* | *K<sub>d</sub>*<br>(μM) | *SEM* | *F<sub>block</sub>* | *SEM* |
| --- | --- | --- | --- | --- | --- | --- | --- | --- |
| **WT-hNav1.7 x WT-KIIIA (5)** | 0.054 | 0.011 | 0.003 | 0.001 | 0.059 | 0.007 | 0.95<sup>a</sup> | 0.033 |
| **Y362C x WT-KIIIA (3)** | 0.101 | 0.016 | 0.044 | 0.004 | 0.436 | 0.052 | 0.75 | 0.055 |
| *N356A x WT-KIIIA (2)* | 0.102 | 0.010 | n.d.<sup>b</sup> | - | n.d. | - | 0.80 | 0.004 |
| *E919Q x WT-KIIIA (3)* | 0.040 | 0.003 | 0.101 | 0.003 | 2.51 | 0.16 | 0.92 | 0.028 |
| *D923A x WT-KIIIA (3)* | 0.083 | 0.022 | 0.193 | 0.022 | 2.34 | 0.41 | 0.89 | 0.037 |

*a* – Fractional block was determined from the Hill fit of concentration-response data (Figure 2B)
*b* – No dissociation was observed during washout, $F_{Block}$ is observed block.
*n* – number of cells tested.

We performed an alanine scan of KIIIA residues that are positioned at the interface with hNa_V_1.7 in our model to determine their effect on binding affinity to the wild-type channel. As with WT-KIIIA, *k_on_*, *k_off_*, and *K_d_* were determined from single exponential fits of peak current during depolarizing voltage steps and are summarized in Table 1; representative data are shown in Figure 2—figure supplement 1A. In the absence of Hill fits of concentration response curves for KIIIA variants, *k_on_* was calculated according to equation 2 (see Materials and Methods). KIIIA substitutions K7A and H12A both had nearly 100-fold decreases in affinity for the wild-type hNa_V_1.7 channel, in agreement with previously published data (Zhang et al., 2007;Zhang et al., 2009;McArthur et al., 2011). We found that KIIIA substitutions W8A and D11A had a 50- and 10-fold reduction in affinity for the wild-type hNa_V_1.7 channel, respectively (Figure 2D). The KIIIA point mutations had little effect on the association rate relative to wild-type KIIIA, with D11A showing the largest effect at a 3-fold increase in association rate (Figure 2D). The observed change in affinity from neutralizing mutations of charged residues was largely driven by 36-fold to 116-fold increases in toxin dissociation rates (Figure 2D). The KIIIA-D11A substitution resulted in both an increase in *k_on_* and *k_off_*. This substitution also would eliminate contact with Na_V_1.7-specific T1398 on the P2-helix in DIII observed in our model. Prior studies had observed that the D11A substitution had no effect on dissociation from rNa_V_1.2, but had a small effect on rNa_V_1.4 binding—slowing *k_off_* 4-fold and very little effect on *k_on_* (Zhang et al., 2007). The reductions in KIIIA affinity from alanine mutations seen *in vitro* correspond to key residues forming the KIIIA-hNa_V_1.7 interface observed in our model (Figure 1B).

We estimated maximal block from fractional current remaining at sub-saturating concentrations of toxin variants by assuming a single binding site with *K_d_* = *k_off_* / *k_on_* and extrapolating to maximal block (see Materials and Methods equations 2, 5, and 7). We caution that these estimates of maximal block are model-dependent and of limited precision, and not as definitive as single channel measurements or experiments in saturating doses of toxin would be. Bearing these cautions in mind, we note that alanine substitution at position 7 (Figure 2C, Table 1) reduces maximal block to 74±4.9%, while substitutions at positions 8 and 11 did not detectably reduce maximal block (90±9.4%, and 100±1.2%, respectively) (Table 1). In addition to levels of block previously reported for K7A, H12A, R10A, R14A variants (McArthur et al., 2011), these results are consistent with an orientation placing K7 towards the acidic residues of the selectivity filter (Figure 1F).

Fractional block at saturating concentrations determined from extrapolation from kinetic data.

### Functional mapping of hNa_V_1.7 residues at the toxin – channel interface support KIIIA binding to the P2-helices in DI and DII

To test the accuracy of the orientation of KIIIA in our model, we used mutations in the P2-helices of DI and DII in the outer pore (Figure 1). We mutated the hNa_V_1.7 N365 and Y362 residues on the P2-helix in DI and E919 and D923 on the P2-helix in DII (Figure 1F). N365A slowed dissociation such that *k_off_* was not measurable during the course of our wash-out experiments, precluding measurement of affinity, but suggesting a limited contribution to toxin binding of this position (Figure 2–Supplement 1, Table 2). Y362C however, produced a modest increase in both association and dissociation yielding a 7.5-fold reduction in affinity corresponding to 1.2 kcal•mol^-1^ (Figure 2E). The E919A mutation did not produce measurable current, yet the E919Q mutation produced functional currents and reduced binding of the wild-type KIIIA by 42-fold corresponding to a 2.2 kcal•mol^-1^ reduction in affinity (Figure 2E). Our model shows a salt bridge between E919 and toxin residue K7 (Figure 1F); the effect of this charge neutralization demonstrates the importance of an acidic residue at this position for toxin binding. The lack of current in the E919A mutant points to a potential steric contribution of this location for pore stability given the proximity to the selectivity filter (Figure 1C, 1F), or poor expression. Our model also shows a hydrogen bond between toxin residue H12 and D923, one helix turn up the DII-P2 helix from E919 (Figure 1F). The substitution D923A likewise reduced affinity of the wild-type KIIIA by 40-fold, or 2.2 kcal•mol^-1^ (Figure 2E) suggesting the elimination of such an interaction. Overall, hNa_V_1.7 mutations Y362C, E919Q, and D923A reduced the binding of the wild-type KIIIA to hNa_V_1.7 in agreement with our structural model of the KIIIA – hNa_V_1.7 complex and published KIIIA – hNa_V_1.2 structure (Pan et al., 2019). The effects of mutations E919Q and D923A in the P2-helix in DII are consistent with the toxin – channel protein-protein interface suggested by our model (see representative data in Figure 2—figure supplement 1B). Overall, our KIIIA alanine scan experiments support the toxin – channel protein-protein interface observed in our KIIIA – hNa_V_1.7 model and the published KIIIA - Na_V_1.2 structure (Pan et al., 2019).

Fractional block at saturating concentrations determined from extrapolation from kinetic data.

### Double Mutant Cycle Analysis confirms pairwise interactions between KIIIA and hNa_V_1.7-DII

Our alanine scan and previous studies have demonstrated the importance of several toxin and channel residues for the binding of KIIIA to hNav1.7. To further validate specific pairwise toxin – channel contacts predicted by our KIIIA – hNa_V_1.7 model, we performed double-mutant cycle analysis experiments (Hidalgo and MacKinnon, 1995;Schreiber and Fersht, 1995;Ranganathan et al., 1996) assessing the coupling between two channel-toxin residue pairs observed in our model, and one pair that was not observed to interact. Specifically, we compared the effects on toxin affinity of hNav1.7 single point mutations and KIIIA single substitution variants at positions corresponding to predicted contact residues, and “double-mutants” consisting of both channel mutations and toxin substitutions (Figure 3A and Figure 3—figure supplement 1). Pairwise contacts can be identified from such experiments on the basis of the path-independence from the wild-type condition to the double-mutant condition: the reduction in binding affinity resulting from a mutation to either side of an interacting pair should be non-additive in the double-mutant condition (Hidalgo and MacKinnon, 1995;Ranganathan et al., 1996). Residue pairs that exhibit additive effects of the double-mutant relative to the single mutants do not make functional interactions contributing to binding. These effects are quantified by calculating the coupling coefficient Ω (Materials and Methods equation 8). Coupling coefficients differing from 1 indicate higher degrees of coupling and are expected with close-contact interactions between the native sidechains (Hidalgo and MacKinnon, 1995;Schreiber and Fersht, 1995;Ranganathan et al., 1996). Importantly, while directly interacting pairs are expected to show coupling, coupling can also result from allosteric effects. We tested the following pairs of double mutants: Nav1.7-E919Q x KIIIA-K7A, D923A x H12A (Figure 3—figure supplement 1), which both interact directly in our model (Figure 4A), and E919Q x D11A, which do not interact directly in our model, with the hypothesis that only the interacting pairs will exhibit coupling coefficients divergent from 1. E919Q x D11A reduced the toxin affinity 240-fold compared to wild-type channel and toxin (*K_d_* =14.2±5.8 μM), and 6- and 21-fold relative to the single mutations, E919Q and D11A (2.51±0.16 μM and 0.66±0.06 μM, respectively) (Figure 3A and Table 3). The large discrepancy between the relative reductions in affinity of the double-mutant compared to wild-type or the single mutants corresponded to a coupling coefficient near 1 (Ω = 0.50±0.38), suggesting little functional interaction between E919 and D11, consistent with the separation of these residues in our model (Figure 3D and Table 4) (Schreiber and Fersht, 1995;Ranganathan et al., 1996). In contrast, the degree to which affinity was weakened by mutations E919Q and K7A together (2.47±1.32 μM) was similar to E919Q (2.51±0.16 μM) or K7A alone (4.29±1.51 μM) (Figure 3A and Table 3), suggesting interaction between E919Q and K7A that is eliminated upon mutation of either side (Ω = 0.014±0.012) (Figure 3D and Table 4). Likewise, the degree to which affinity was weakened by mutations D923A and H12A together (2.40±1.45 μM) was similar to D923A (2.34±0.41 μM) or H12A alone (7.41±4.6 μM) (Figure 3A and Table 3). The non-additive effect of these substitutions suggests close contact between D923 and H12 is eliminated upon mutation of either side (Ω = 0.0082±0.0091) (Figure 3D and Table 4). The coupling observed between these pairs are consistent with pairwise interactions between charged amino acids (Hidalgo and MacKinnon, 1995;Schreiber and Fersht, 1995;Ranganathan et al., 1996). These results are consistent with E919 – K7 and D923 – H12 pairwise interactions observed in both our model (Figure 4A) and the recent structure of KIIIA - Na_V_1.2 complex (Pan et al., 2019), which both show strong electrostatic interactions between these residue pairs, providing further experimental validation of the KIIIA binding pose observed in our model.

**Figure 3.**
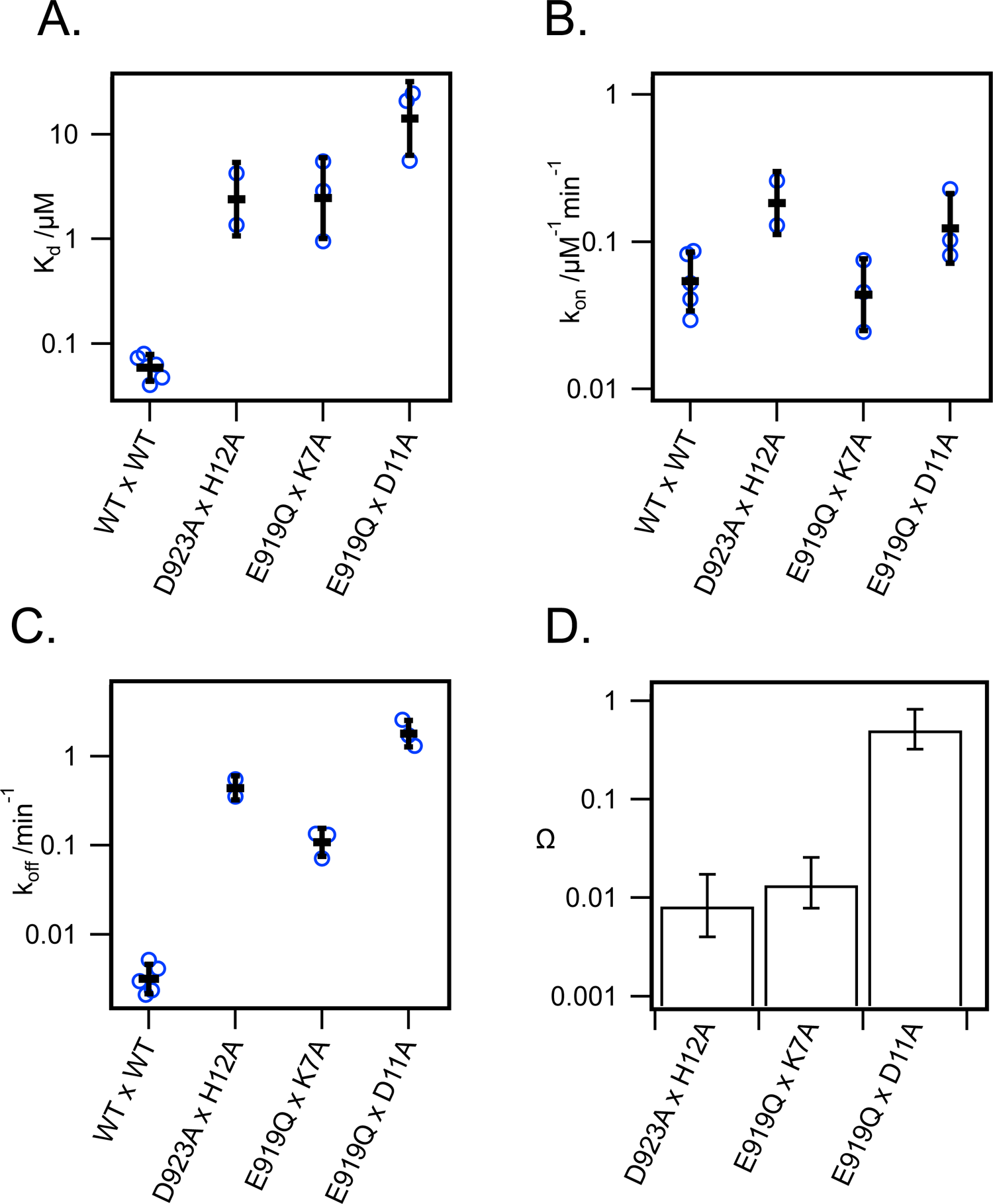
Double mutant cycle analysis supports key pairwise interactions between KIIIA and hNa_V_1.7. (**A**) *K_d_*, (B) *k_on_*, and (C) *k_off_* from double-mutant experiments of thermodynamic double-mutant cycles between H12A x D923A, K7A x E919Q, and D11A x E919Q. Bars are geometric mean±SD, n=2-5 cells per condition (reported in table 3), and empty circles are individual cells. (D) Coupling coefficients Ω±error calculated from linear propagation of SEM of *K_d_* values from single and double-mutation experiments reported in Table 3.

**Table 3.**
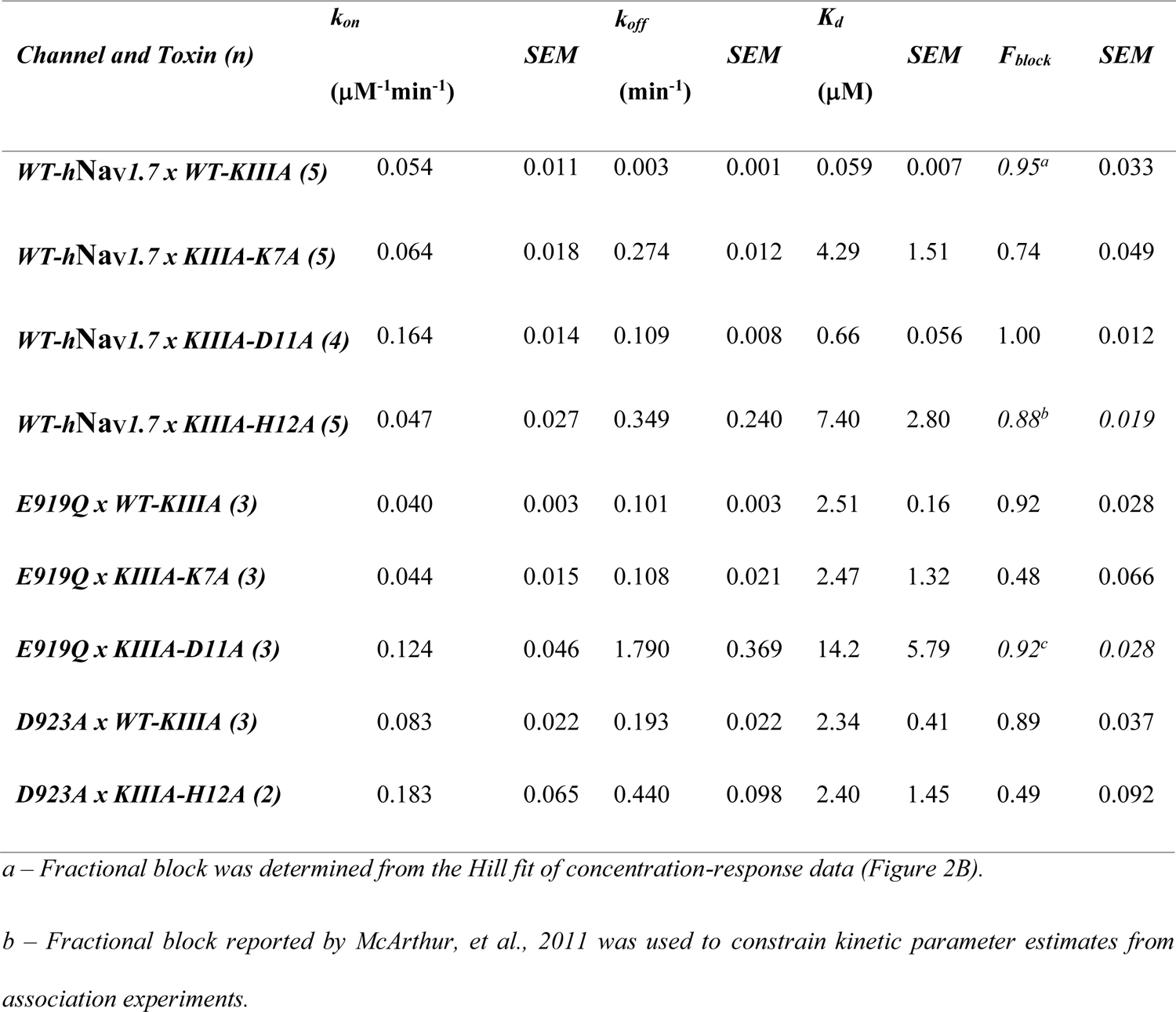
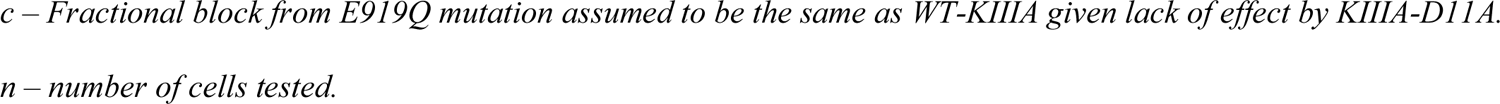
Block of double-mutant cycle pairs of hNa_V_1.7 and KIIIA

| *Channel and Toxin (n)* | $k_{on}$<br>($\mu\text{M}^{-1}\text{min}^{-1}$) | *SEM* | $k_{off}$<br>($\text{min}^{-1}$) | *SEM* | $K_d$<br>($\mu\text{M}$) | *SEM* | $F_{block}$ | *SEM* |
| --- | --- | --- | --- | --- | --- | --- | --- | --- |
| **WT-hNav1.7 x WT-KIIIA (5)** | 0.054 | 0.011 | 0.003 | 0.001 | 0.059 | 0.007 | 0.95<sup>a</sup> | 0.033 |
| **WT-hNav1.7 x KIIIA-K7A (5)** | 0.064 | 0.018 | 0.274 | 0.012 | 4.29 | 1.51 | 0.74 | 0.049 |
| **WT-hNav1.7 x KIIIA-D11A (4)** | 0.164 | 0.014 | 0.109 | 0.008 | 0.66 | 0.056 | 1.00 | 0.012 |
| **WT-hNav1.7 x KIIIA-H12A (5)** | 0.047 | 0.027 | 0.349 | 0.240 | 7.40 | 2.80 | 0.88<sup>b</sup> | 0.019 |
| **E919Q x WT-KIIIA (3)** | 0.040 | 0.003 | 0.101 | 0.003 | 2.51 | 0.16 | 0.92 | 0.028 |
| **E919Q x KIIIA-K7A (3)** | 0.044 | 0.015 | 0.108 | 0.021 | 2.47 | 1.32 | 0.48 | 0.066 |
| **E919Q x KIIIA-D11A (3)** | 0.124 | 0.046 | 1.790 | 0.369 | 14.2 | 5.79 | 0.92<sup>c</sup> | 0.028 |
| **D923A x WT-KIIIA (3)** | 0.083 | 0.022 | 0.193 | 0.022 | 2.34 | 0.41 | 0.89 | 0.037 |
| **D923A x KIIIA-H12A (2)** | 0.183 | 0.065 | 0.440 | 0.098 | 2.40 | 1.45 | 0.49 | 0.092 |

<sup>a</sup> – Fractional block was determined from the Hill fit of concentration-response data (Figure 2B).
<sup>b</sup> – Fractional block reported by McArthur, et al., 2011 was used to constrain kinetic parameter estimates from association experiments.
*c* – Fractional block from E919Q mutation assumed to be the same as WT-KIIIA given lack of effect by KIIIA-D11A. *n* – number of cells tested.

**Table 4.** Coupling coefficients from double-mutant cycle experiments

|  | *D923A x H12A* | *error* | *E919Q x K7A* | *error* | *E919Q x D11A* | *error* |
| --- | --- | --- | --- | --- | --- | --- |
| **Ω** | 0.0082 | 0.0091 | 0.014 | 0.012 | 0.50 | 0.38 |

*Error calculated by linear propagation of uncertainty (Ku, 1966;Hidalgo and MacKinnon, 1995).*

### Na_V_ channel isoforms with distinct toxin affinities have divergent residues at the KIIIA binding interface

The differences in KIIIA binding affinity between the Na_V_ channel isoforms likely arise from variations in sequence in the P2 helices and extracellular loop regions (Figure 4A and B). The published structure of the KIIIA - hNa_V_1.2 complex (Pan et al., 2019) was not available when we generated our structural model of the KIIIA - hNa_V_1.7 complex, yet it is consistent with our structural model and further supports this observation. The backbone root mean square deviation (RMSD) of the KIIIA - hNa_V_1.7 model and the KIIIA - hNa_V_1.2 structure over KIIIA and the P2-helices is ∼1.0 Å (Figure 1—figure supplement 1B). The specific pairwise contacts between K7-E919 and H12-D923 in our KIIIA - hNa_V_1.7 model, validated by our mutant cycle analysis, are in agreement with the corresponding pairwise contacts between K7-E945 and H12-D949 observed in the KIIIA - hNa_V_1.2 complex structure (Figure 4A and B) (Pan et al., 2019). Our KIIIA - hNa_V_1.7 model also predicted other contacts observed in the published KIIIA - hNa_V_1.2 structure (Pan et al., 2019), including pairwise interactions between KIIIA N3 and W8 with E307 and Y339, respectively, on the extracellular S5-P1 loop in DI (Figure 4A and B).

The KIIIA critical residues D11 and R10 in our KIIIA - hNa_V_1.7 model are positioned similarly in the KIIIA - hNa_V_1.2 structure, but details of toxin – channel interactions involving these residues are different (Figure 4A). D11 forms a hydrogen bond with T1398 on the P2-helix in DIII in our KIIIA - hNa_V_1.7 model (Figure 4A), but the substitution of Thr (T1398) hNa_V_1.7 to Met (M1425) on the P2-helix in DIII in hNa_V_1.2 removes this interaction, and a new hydrogen bond is formed between D11 with the nearby residue Y1429 (Figure 4—figure supplement 1) (Pan et al., 2019). In the hNa_V_1.2 structure, R10 interacts with D1426 on the P2-helix in DII, but the corresponding position in hNa_V_1.7 is I1399, eliminating possible charge interaction with this residue in hNa_V_1.7 (Figure 4A and B). This difference potentially contributes to the R10 interaction with the nearby acidic residue D1662 on the extracellular S5-P1 loop in DIV of hNa_V_1.7 (Figure 4—figure supplement 1). It is noticeable that Asp at position 1662 is unique to hNa_V_1.7 - corresponding residues at this position in other Na_V_ channel subtypes are Val, Ala, and Ser (Figure 4B). Additionally, the corresponding residues to T1398 and I1399 on the P2-helix in DIII of hNa_V_1.7 are Met and Asp, respectively, in all other human Na_V_ channels. Because of that, it seems reasonable to hypothesize that these residues provide a major contribution to structural determinant of KIIIA interaction with hNav1.7.

The R14 residue on KIIIA is important for KIIIA binding to Nav channels and the substitution of Arginine to Alanine at this position has been shown to gain selectivity for hNa_V_1.7 versus hNa_V_1.2 (McArthur et al., 2011). Both our KIIIA - hNa_V_1.7 model and the KIIIA - hNa_V_1.2 structure show the positively charged R14 forming cation – π interactions with Y1416 (hNa_V_1.7) or Y1443 (hNa_V_1.2) on the extracellular P2-S6 loop in DIII (Figure 4A). Notably, in the KIIIA - hNa_V_1.2 structure R14 is also in proximity to the negatively charged E919 (hNa_V_1.2 numbering) on the extracellular S5-P1 loop in DII (Pan et al., 2019). However, in our KIIIA - hNa_V_1.7 model, R14 is in proximity to T893 on the extracellular S5-P1 loop in DII (which is corresponding to E919 in hNa_V_1.2) and E1417 on the extracellular P2-S6 loop in DIII (Figure 4A and B). We reason that these differences may contribute to the reported selectivity of KIIIA R14A substitution against hNav1.7. Overall, despite the similarity in binding pose and channel architecture, sequence variance at KIIIA binding site, comprised of the P2 helices and extracellular loop regions likely contribute to differences in KIIIA binding affinity of between hNav1.7 and hNav1.2, and perhaps also among other channel isoforms.

**Figure 4.**
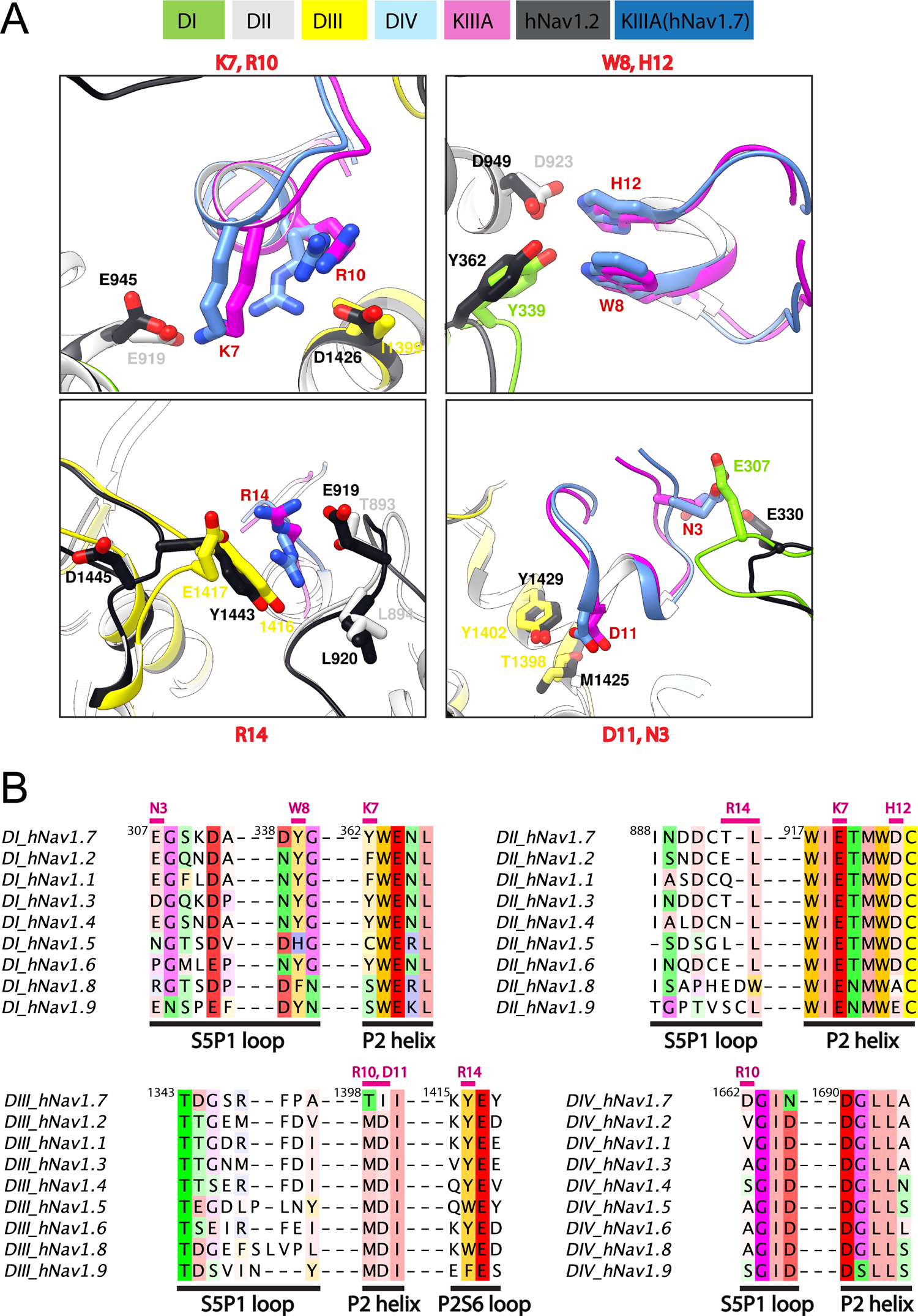
Marked difference for KIIIA binding specificity among Na_V_ channel isoforms. (**A**) Comparisons of key residue interactions on the KIIIA - hNa_V_1.7 model and the KIIIA - hNa_V_1.2 structure. Domains and residue labels are depicted according to color keys. (**B**) Sequence alignment of the different Na_V_ subtypes at the interaction site with KIIIA. Different regions are labeled. Key interacting residues of KIIIA are labeled and colored in magenta at positions of corresponding interactions in channel residues. Superscripts show sequence number of hNa_V_1.7.

### Structural dynamics of KIIIA binding to hNa_V_1.7 and hNa_V_1.2 revealed by molecular dynamics simulations

To further study the molecular mechanism of the KIIIA interaction with hNa_V_1.7, we performed molecular dynamics (MD) simulations of our KIIIA - hNa_V_1.7 complex model. The 3 independent 1.5 μs MD simulations revealed highly dynamic KIIIA binding with the backbone RMSDs between KIIIA and hNa_V_1.7 interface fluctuating between 3.5 – 4.5 Å (Figure 5 – figure supplement 1). Remarkably, we observed key positively charged residues K7 and R10 on KIIIA interacted not only with the acidic residue on the P2-helix in DII identified from our model (E919 and D923) but also with acidic residues on the P2-helix in DIV. The density projection of these residues on the membrane plane revealed the primary amine group of K7 was mostly localized near D923 and E919 on P2-helix in DII and D1690 on P2-helix in DIV (see sites A1, A2 and A3 in Figure 5A) while R10 was localized near E919 (DII) and D1690 (DIV) (Figure 5 – figure supplement 2). The dynamic interactions of K7 with multiple acidic residues on P2 helices is consistent with our functional characterization of KIIIA - hNa_V_1.7 interactions where we observed a 100-fold reduction in *K*_d_ for the K7A mutation on KIIIA and only a 42-fold reduction for E919Q mutation on the channel. As an indication of highly dynamic interactions of KIIIA with hNav1.7, we also observed key residues on KIIIA formed interactions with multiple residues on the channel as shown in the fractional contact map (Figure 5). Notably, W8 frequently formed contacts with K310 and Y339 on the extracellular S5-P1 loop in DI. H12 interacted mainly with D923 on the P2-helix in DII and also with the backbone of P895 on the extracellular S5-P1 loop in DII. D11 is positioned deep at the interface between DII and DIII within a region formed by hNa_V_1.7 residues R911 on the P1-helix in DII, E919 on the P2-helix in DII, and Y1402 on the P2-helix in DIII. R14 primarily interacted with Y1416 on the extracellular P2-S6 loop in DIII, and T893, L894, P895 and R896 on the extracellular loop in DII and did not maintain interaction with E1417 on the extracellular loop in DIII as initially identified in our model (Figure 5—figure supplement 2). In addition, during equilibration we consistently observed the initial placed sodium ion in the selectivity filter localized near E916 and E919 on the P1-P2-helix region in DII (Figure 5A – figure supplement 1), which agrees with the density identified as a sodium ion at the same position in the structure of the KIIIA - hNa_V_1.2 complex (Pan et al., 2019). This sodium ion quickly diffused out to the extracellular bulk via an open passage formed between KIIIA and the selectivity filter region of the channel. The sodium ion escape is consistent with the incomplete block of Na_V_ current observed in experiments (Figure 2) and incomplete reduction of the unitary Na_V_ channel conductance when KIIIA is bound (Zhang et al., 2007;McArthur et al., 2011).

**Figure 5.**
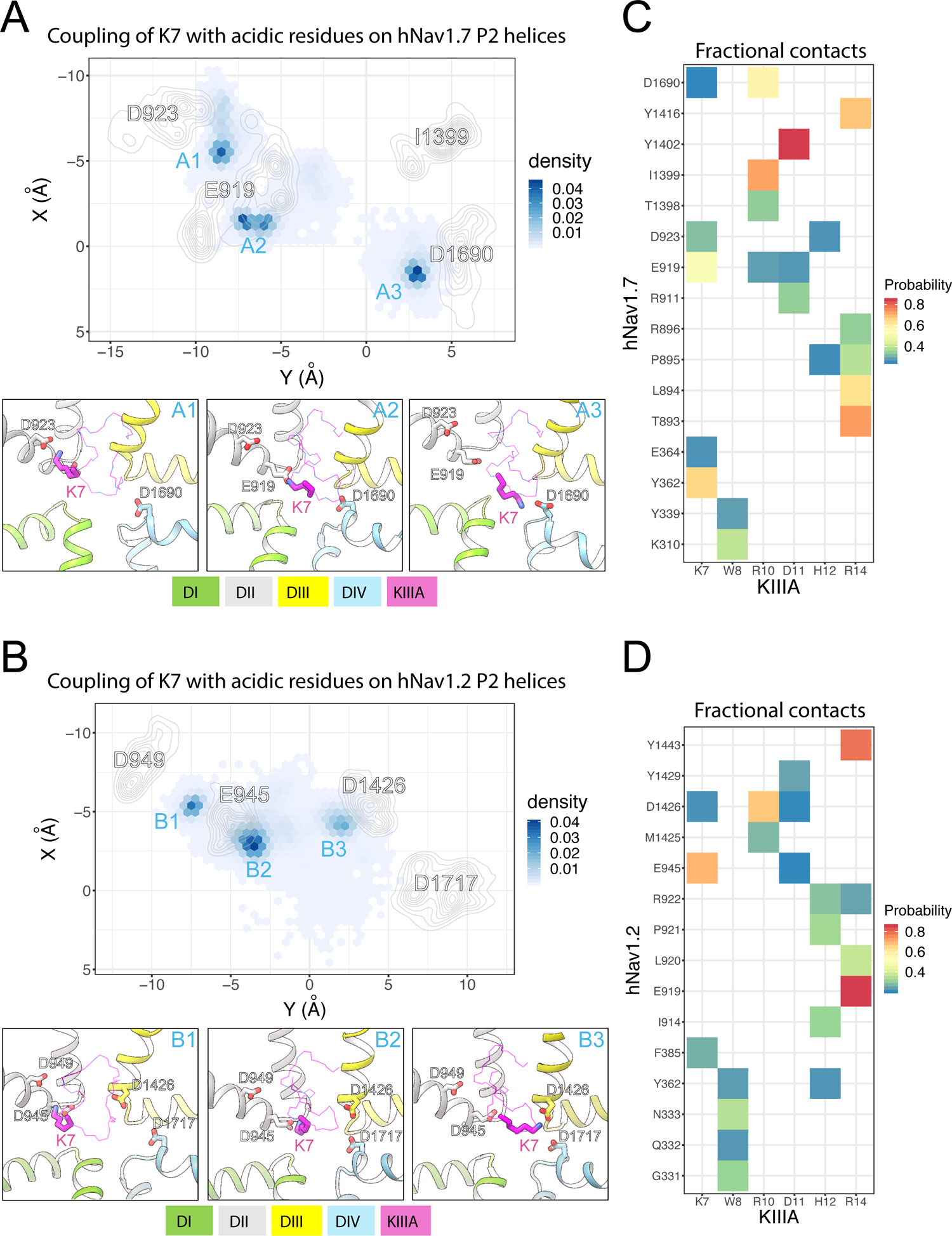
MD simulations of KIIIA - hNa_V_1.7 and KIIIA - hNa_V_1.2 complexes show structural dynamics of KIIIA interaction with hNa_V_1.7 (A and C) and hNa_V_1.2 (B and D). (A, B) Couplings of K7 with key acidic residues on the hNa_V_ P2 helices. Density projections of K7 (Nitrogen atom) and the acidic residue (oxygen atoms) on P2-helix (labeled) on the XY plane (upper panel). For clarity, densities of the acidic residues are shown as contour plots using kernel density estimation. Representative snapshots showing interactions of K7 with the acidic residues at high density regions (labeled) identified from the density projections (lower panel). (C, D) Heatmaps showing fractional contacts between key residues on KIIIA and hNa_V_ channels during the simulations.

We used the cryoEM structure of KIIIA - hNa_V_1.2 complex (Pan et al., 2019) to study structural dynamics at the toxin – channel interface and compare them to dynamics observed in our KIIIA - hNa_V_1.7 complex model (Figure 5A). We conducted 3 independent 1 μs MD simulations using the same procedure as applied for the KIIIA - hNa_V_1.7 model (see Materials and Methods). Similar to the simulations of our KIIIA – hNav1.7 model, the KIIIA - hNa_V_1.2 complex structure was dynamic during simulations with the RMSDs at the toxin – channel interface between 3 - 5 Å (Figure 5 – supplement 1). We also observed that key positively charged residues K7 and R10 on KIIIA interacted with multiple acidic residues on the P2 helices of DI, DII, and DIV on hNa_V_1.2 (Figure 5B). The primary amine group of K7 was mainly localized near E945 and D949 on the P2-helix in DII and D1426 on the P2-helix in DIII (see sites B1, B2 and B3 in Figure 5B). Interactions with the conserved acidic residues D1717 on the P2-helix in DIV (corresponding to D923 and D1690 in hNa_V_1.7, respectively) were more limited compared to that in the simulations of KIIIA - hNa_V_1.7 model (see site A3 in Figure 5A). Similarly, R10 was localized near E945 on the P2- helix in DII and D1426 on the P2-helix in DIII while interactions with the conserved acidic residue D1717 (D1690 in hNa_V_1.7) were less dominant compare to that in hNa_V_1.7 (Figure 5 – figure supplement 1). These differences between hNa_V_1.2 and hNa_V_1.7 can be explained by a contribution of a single residue substitution in P2-helix in DIII. Particularly, D1426 on the P2-helix in DIII of hNa_V_1.2 is conserved among all human Na_V_ channel isoforms, except for hNa_V_1.7, which has I1399 at this position (Figure 4). The absence of an acidic residue, Asp at the 1399 position in hNa_V_1.7 (D1426 in hNa_V_1.2), substituted by a hydrophobic residue, I1399 may facilitate interactions of positively charged K7 and R10 with the nearby acidic residue D1690 on the P2-helix in DIV in hNa_V_1.7 (Figure 5—figure supplement 1). Indeed, functional studies showed that the mutation R10A produced a 35% reduction in KIIIA block of rNa_V_1.4, while the loss of toxin block caused by the R10A mutation was largely rescued by hNa_V_1.7-like Ile on the P2-helix in DIII (McArthur et al., 2011). Based on these observations, we speculate that dynamic interactions of basic residues K7 and R10 on KIIIA with acidic residues on P2-helix on hNa_V_ channels, driven by sequence variability at toxin – channel interface may provide a structural basis for differences in KIIIA interactions among hNa_V_ channel isoforms. The sodium ion in the KIIIA - hNa_V_1.2 structure at the site formed by E942 and E945 (corresponding to E916 and E919 in hNa_V_1.7) on the P1-P2-helix region in DII of hNa_V_1.2 quickly diffused out to the extracellular bulk, in agreement with an incomplete block by KIIIA (Zhang et al., 2007) and our simulation of the KIIIA - hNa_V_1.7 model (Figure 5A and B). We did not observe re-entering events of sodium ions into the hNa_V_1.2 selectivity filter, similarly to our simulations of the KIIIA - hNa_V_1.7 model.

## Discussion

### Computational modeling and functional testing reveal molecular determinants of μ-conotoxin KIIIA interaction with Na_V_1.7

Our computational model of hNa_V_1.7, docking of KIIIA to the hNa_V_1.7 model, and functional testing of KIIIA and hNa_V_1.7 mutations presented here were completed prior to the publication of KIIIA - hNa_V_1.2 complex structure (Pan et al., 2019). While hNa_V_1.7 and hNa_V_1.2 channels share sequence homology within most of the KIIIA binding region, they exhibit several key differences at the toxin - channel interface. We functionally characterized specific pairwise contacts between KIIIA and hNa_V_1.7 predicted by our model, particularly between K7 on KIIIA and E919 on the P2-helix in DII and also between H12 on KIIIA and D923 on the P2-helix in DII. These pairwise interactions agree with the corresponding pairwise interactions between K7-E945 and H12-D949 observed in the KIIIA - hNa_V_1.2 complex structure (Figure 4A and B) (32). Interestingly, we also observed that our KIIIA – hNa_V_1.7 complex model based on the eeNa_V_1.4 structure is similar to our earlier KIIIA – hNa_V_1.7 complex model based on the bacterial Na_V_Ab channel structure using experimental restraints (PDB: 3RVY (Payandeh et al., 2011)). In this effort, the pairwise distance constraints from available experimental data were used to model the interactions of GIIIA toxin, a close homolog of KIIIA with the homology model of hNav1.4 based on the bacterial NavAb structure. The model GIIIA – hNav1.4 were then subsequently used to build the KIIIA – hNav1.7 model. It predicted accurately the KIIIA orientation and both the interaction of K7 with E919 and H12 with D923 (Figure 1—figure supplement 1A). However, the NavAb based model failed to identify the pairwise interaction of W8 and Y339 (interacted with E926 instead). This likely due to the limitation of the bacterial template with the difference in pore architecture compared to mammalian Nav channels and the absence of extracellular loop regions in the NavAb based model. Together, our work has demonstrated the predictive power of structural modeling with different degrees of experimental restraints to study peptide toxin – channel interactions.

The Cryo-EM structures of Na_V_1.1, Na_V_1.2, Na_V_1.3, Na_V_1.4, Na_V_1.5, Na_V_1.6, Na_V_1.7, and Na_V_1.8 channels (Pan et al., 2018;Pan et al., 2019;Shen et al., 2019;Jiang et al., 2020;Jiang et al., 2021;Pan et al., 2021;Huang et al., 2022a;Huang et al., 2022b;Li et al., 2022;Fan et al., 2023) revealed that the extracellular vestibule of the channel pore targeted by KIIIA is surrounded by several relatively long loop regions, raising the possibility that the access pathway to the KIIIA binding site is relatively narrow. Restricted access and escape pathways for KIIIA binding could confer the relatively slow *k_on_* and *k_off_* rates observed in our functional studies (Figures 2 and 3) and previously published data (Zhang et al., 2007;McArthur et al., 2011;Wilson et al., 2011). We speculate that variability in the Na_V_ channel extracellular vestibule loops could underlie the differences in KIIIA kinetics.

While our KIIIA - hNa_V_1.7 model and the published KIIIA - hNa_V_1.2 structure are supported by functional data, further interactions of key charged residues K7 and R10 on KIIIA with multiple acidic residues on the channels was observed in our MD simulations and could provide insight into selectivity of the toxin and channel block. Notably, the unique residue I1399 on the P2-helix in DIII of hNa_V_1.7 may contribute to KIIIA binding on hNa_V_1.7 by influencing the interactions of K7 and R10 on KIIIA with acidic residues on the channel. We also observed escapes of sodium ions initially located in the selectivity filter in both KIIIA - hNa_V_1.7 and KIIIA - hNa_V_1.2 which agrees with functional data showing that KIIIA incompletely blocks Na_V_ channels. Interestingly, despite having 150 mM of NaCl in the bulk solvent, we did not observe re-entering events of sodium ions in the selectivity filter during the 3 independent 1.5 μs MD simulations. This possibly can be explained by the relatively high percentage (> 94%) of KIIIA block observed in both hNa_V_1.7 and hNa_V_1.2, which implied such re-binding events are rare and may not be observed in our conventional 1.5 μs MD simulation trajectories. Our results agree with the functional data showing that KIIIA is a highly efficient but incomplete pore blocker of Na_V_ channels. Study of the full binding and blocking mechanism of KIIIA on Na_V_ channels is beyond the scope of this study and will require much longer simulation times or enhanced sampling techniques, and a structure of KIIIA bound to an open and conductive state of Na_V_ channel, which is not currently available. Our MD simulations results reflect dynamics of the binding configuration captured in our KIIIA - hNa_V_1.7 model and the KIIIA - hNa_V_1.2 structure to provide a more comprehensive insights into KIIIA – Nav channel interactions that extend beyond interpretation from the static structures.

We used an integrative Rosetta computational modeling, functional characterization, and MD simulations approach to study molecular determinants of peptide toxin – Nav channel interactions. Establishing successful predictions with computational modeling is a critical step towards computational design of selective and potent peptide-based therapeutics. Our approach can be potentially expanded to rational design of novel peptides to target the extracellular pore vestibule region of Na_V_ channels. Despite the high sequence conservation in the pore region of Na_V_ channels, our work shows that specific sequence differences between Na_V_ channels in the extracellular loop regions and the P2-helices of the pore can have important functional consequences on toxin – channel interaction. Rosetta protein design (Kuhlman et al., 2003;Cao et al., 2020;Leman et al., 2020) and optimization (Silva et al., 2019;Linsky et al., 2020) informed by these structural insights could potentially lead to development of high-affinity and specificity peptide inhibitors of Na_V_ channels (Nguyen et al., 2022;Nguyen and Yarov-Yarovoy, 2022), forming a new class of biologics to treat Na_V_ channel related diseases.

## Conclusions

We generated a structural model of the conotoxin KIIIA in complex with hNa_V_1.7 using homology modeling and docking simulations. Our model was validated with functional testing, using alanine-scan mutagenesis of KIIIA and hNa_V_1.7, double mutant cycle analysis of specific pairwise toxin – channel interactions, supporting that acidic residues E919 and D923 on the P2-helix in DII of Na_V_1.7 significantly contribute to toxin – channel interaction, and that KIIIA forms multiple interactions with the extracellular loops in DI-III. The published structure of the KIIIA - hNa_V_1.2 complex further supports predictions observed in our model. Unbiased MD simulations of KIIIA - hNa_V_1.7 and KIIIA - hNa_V_1.2 complexes suggest a potential important role of I1399 on the P2 helix in DIII of hNa_V_1.7 that may underlie the structural basis of KIIIA block of hNa_V_1.7 conductance. Overall, our results further characterize the molecular determinants of the KIIIA interaction with human Nav channels and can be potentially useful for rational design of increases in the potency or relative selectivity towards Nav1.7. As Nav1.7 is a drug target for pain (Payandeh and Hackos, 2018;Bennett et al., 2019;Dib-Hajj and Waxman, 2019), such optimization of KIIIA could result in novel peptide-based therapeutics.

## Acknowledgments

We would like to thank Drs. Heike Wulff, Jie Zheng, and Igor Vorobyov, and members of Yarov-Yarovoy and Sack laboratories for helpful discussions. We thank Dr. Nieng Yan (Princeton University) for independent comparison of our KIIIA – hNa_V_1.7 model to coordinates of KIIIA – hNa_V_1.2 structure (Pan et al., 2019) prior to release in Protein Data Bank, and sharing the coordinates of electric eel and human Na_V_1.2 channel structures. We thank Dr. Christoph Lossin (University of California, Davis) for providing the hNa_V_1.7 cell line and WT-hNa_V_1.7 channel construct, as well as Dr. William Catterall (University of Washington) for the tsa201 cell line. Anton 2 computer time was provided by the Pittsburgh Supercomputing Center (PSC) through Grant R01GM116961 from the National Institutes of Health. The Anton 2 machine (Shaw et al., 2014) at PSC was generously made available by D.E. Shaw Research. This research was supported by U.S. National Institutes of Health awards U01HL126273 and R01HL128537 to VYY, R01NS096317 to JTS, UC Davis Academic Senate Award FL18YAR to VYY, NIH T32 GM099608 to IHK, and AHA 17PRE33670204 to IHK.

## Materials and methods

### Homology modeling of hNa_V_1.7 based on EeNa_V_1.4 structure

The cryo-EM structure of the Na_V_1.4-β1 complex from the electric eel (eeNa_V_1.4) (PDB ID: 5XSY) (Yan et al., 2017) was used to generate the model of hNa_V_1.7 channel using Rosetta structural modeling software (Song et al., 2013;Bender et al., 2016). Initially, we refined the published coordinates of eeNa_V_1.4, without the β1 subunit by using the Rosetta cryo-EM refinement protocol (Wang et al., 2016) and picked the lowest scoring density-refitted eeNa_V_1.4 model to use as a template. The comparative modeling protocol RosettaCM (Song et al., 2013) was then used in combination with the electron density of the eeNa_V_1.4 to model the hNa_V_1.7 structure. We generated 5,000 structural models of hNa_V_1.7 and selected the top 500 lowest-scoring models for clustering analysis as described previously (Bonneau et al., 2002). Visual inspection of the top scoring clustered models was used to select the final model for the docking study.

### Molecular docking of KIIIA to the hNa_V_1.7 model

The solution NMR structure of KIIIA (PDB ID: 2LXG) (Khoo et al., 2012) was used as an ensemble to dock to the hNa_V_1.7 model using Rosetta protein-protein docking approach (Fleishman et al., 2011;Bender et al., 2016). By default, Rosetta moves proteins apart at the beginning of the protein-protein docking procedure which led to placement of KIIIA above the extracellular pore loops and did not allow sampling of the KIIIA binding site within the selectivity filter region because the KIIIA was not able to pass the narrow passage created by the extracellular pore loops. To address this problem, we subsequently divided the docking protocol into two stages (see details of Rosetta commands and scripts in Supplementary File 1). In stage 1, docking was performed with the DI S5P1 and DIII S5P1 loops truncated, and full random translational and rotational perturbation of KIIIA at both low and high-resolution phases. This stage generated 20,000 structural models of the docking complexes. We then selected the top 1,000 models based on the total scores and filtered based on the Rosetta ΔΔG (an estimate of the binding energy of a complex) to select the top 500 models. ΔΔG is computed by taking the difference of the energy of the KIIIA – hNa_V_1.7 complex and of the separated KIIIA and hNa_V_1.7 structures. We clustered these complexes using the Rosetta legacy clustering application. The center models of top 20 clusters then passed to stage 2 docking. In this stage, positions of KIIIA in the top 20 clusters were used to create 20 different starting docking trajectories with the full structure of hNa_V_1.7 model including all the extracellular loop regions. The full translational and rotational perturbation used in the previous stage was turn off. Instead, only limited local perturbation was allowed in both centroid and full-atom refinement phases. Similar to stage 1, we generated 20,000 structural docking models and filtered based on the Rosetta total score and ΔΔG to select the top 500 models, which were again clustered to finalize the top 5 complexes for visual inspection. The selected docking model presented here (see coordinates of our KIIIA – hNa_V_1.7 model in Supplement File – Model 1) is the only one in the top 5 clusters models that has KIIIA partially occluding the pore and K7 near the selectivity filter in agreement with experimental data demonstrating the contribution of K7 to binding affinity and percentage block (Zhang et al., 2007;McArthur et al., 2011).

### Molecular dynamics simulation of KIIIA - hNa_V_1.7 and KIIIA - hNa_V_1.2 complexes

The docking complex of KIIIA - hNa_V_1.7 and the cryo-EM structure of hNa_V_1.2 (PDB ID: 6J8E) were used to setup systems for MD simulations. For the hNa_V_1.2 structure, Rosetta density refinement protocol was applied as described above for the hNa_V_1.7. The missing region on DI extracellular loop was modeled using Rosetta loop modeling. We placed one sodium ion in the selectivity filter and one in the cavity of the channels as initial setup for both simulations. CHARMM-GUI (Jo et al., 2008) was used to embed the KIIIA - hNa_V_1.7 model and the KIIIA - hNa_V_1.2 structure (PDB ID: 6J8E) in a lipid bilayer of POPC with explicit TIP3P water molecules at a concentration of 150 mM NaCl. CHARMM36 forcefield was used for proteins, lipids, ions, and waters in both systems. Each system contains approximately of 164,000 atoms. Protonation state is assigned at neutral pH. N-epsilon nitrogen of H12 on KIIIA is protonated instead of N-delta nitrogen as suggested in both the Rosetta model of hNa_V_1.7 – KIIIA and the hNa_V_1.2 – KIIIA structure. The C-terminal of KIIIA is amidated to be consistent with the KIIIA variant used in our experiments.

Equilibrations were run on our local GPU cluster using NAMD version 2.12 (Jiang et al., 2011). After 10,000 steps of steepest descent minimization, MD simulations started with a timestep of 1 fs with harmonic restraints initially applied to protein heavy atoms and some lipid tail dihedral angles as suggested by CHARMM-GUI (Jo et al., 2008). These restraints were slowly released over 2 ns. Harmonic restraints (0.1 kcal/mol/Å^2^) were then applied only to protein backbone atoms, and the systems were equilibrated further for 20 ns with a timestep of 2 fs. All bonds to H atoms were constrained using the SHAKE algorithm in order to use a 2 fs timestep. Simulations were performed in NPT ensemble with semi-isotropic pressure coupling to maintain the correct area per lipid, and constant temperature of 303.15 K. Particle Mesh Ewald (PME) method was used to compute electrostatic interactions. Non-bonded pair lists were updated every 10 steps with a list cutoff distance of 16 Å and a real space cutoff of 12 Å with energy switching starting at 10 Å. Independent simulation systems were created by using different seed numbers in the equilibrations.

We used the Anton 2 software version 1.31.0 for production runs of each system on the Anton 2 supercomputer. We ran 3 independent simulations of 1.5 μs for our hNa_V_1.7 – KIIIA model and 3 independent simulations of 1μs for the hNa_V_1.2 – KIIIA structure. Simulations were performed in the NPT ensemble at 303.15 K A with 2 fs timestep. Non-bonded long-range interactions computed every 6 fs using the RESPA multiple time step algorithm. The multi-integrator algorithm was used for temperature and semi-isotropic pressure coupling and the u-series algorithm was used for long-range electrostatic interactions. A long-range Lennard-Jones (LJ) correction (beyond cutoff) was not used as was suggested for CHARMM36 lipid force field.

### Modeling of KIIIA - hNa_V_1.7 complex using Na_V_Ab structure

We previously performed homology modeling of human Na_V_ channels based on bacterial Na_V_ channel structure before any eukaryotic Na_V_ channel structures were published. We first generated a model of hNa_V_1.4 pore using x-ray structure of the bacterial Nav channel Na_V_Ab (PDB ID: 3RVY) as a template using Rosetta homology modeling (Bender et al., 2016). We selected to first model Na_V_1.4 channel because of availability of experimental data on conotoxin – Na_V_ channel interactions for model validation (Dudley et al., 2000;Choudhary et al., 2007). The extracellular loop regions in hNa_V_1.4 pore model were truncated and the P2 loops were rebuilt *de novo* using the Rosetta homology modeling (Bender et al., 2016). We used available experimental data for conotoxin GIIIA (homolog of KIIIA) interaction with Na_V_1.4 channel (Dudley et al., 2000;Choudhary et al., 2007) to guide the docking of GIIIA to hNa_V_1.4 model. Specifically, charged residues R13 and K16 on GIIIA were biased to interact with acidic residues on the Na_V_1.4 P2 helices during docking using Rosetta bounded restraints (Bender et al., 2016). We then used the top GIIIA - hNav1.4 model that agreed with experimental data (Dudley et al., 2000;Choudhary et al., 2007) to build a model of GIIIA - hNa_V_1.7 complex using Rosetta homology modeling (Bender et al., 2016). To create the initial configuration for KIIIA docking, we superimposed KIIIA onto GIIIA based on GIIIA - hNa_V_1.7 complex model. In this final docking step, KIIIA was docked using only full-atom docking perturbations following by a flexible backbone refinement using Rosetta FlexPepDock (Raveh et al., 2010) and the best model was selected using Rosetta interface score. We did not perform experimental characterization on Na_V_Ab based KIIIA - hNa_V_1.7 complex model because the subsequently published structures of eukaryotic Na_V_ channels have allowed us to perform higher accuracy homology modeling of hNa_V_1.7 based on eeNa_V_1.4 structure (described above).

### Cell culture, transfection, and preparation

Electrophysiology experiments were performed on transiently transfected tsa-201 cells (gift from William Catterall) and a HEK 293T cell line stably expressing hNa_V_1.7 (gift from Chris Lossin). Cells were grown at 37°C, 5% CO_2_ in DMEM with 4.5g/L D-glucose, L-glutamine, and 110 mg/L Sodium Pyruvate (Gibco cat# 11995-065) with 10% FBS, and 100 units/mL Penicillin/Streptomycin (Gibco cat# 15140-122). The stable cell line was raised in the same conditions with 500 μg/mL G418 as a selection agent. Cells were grown to 70% confluency in 35mm dishes and passaged every 2-3 days for tsa-201 and 3-4 days for the stable-cell line. Cells were washed with divalent-free DPBS (Gibco cat# 14190-144) and dissociated with 0.05% Trypsin-EDTA (Gibco cat# 25300-054) and seeded to fresh dishes with pre-warmed media. tsa-201 cells were transfected via Lipofectamine 2000 24-48 hours prior to experiments with 1 μg pCMV6-SCN9A (gift from Dr. Christoph Lossin) and 0.5 μg pMaxGFP (Lonza) for identification of transfected cells. Mutant constructs were purchased, and coding sequences verified by Mutagenex. Prior to experiments, cells were washed with DPBS and dissociated in 1mL Versene (Gibco cat# 15040-066) and scraped from the dishes and transferred to a 14mL conical tube with 3 mL DMEM. They were centrifuged at 1000 x *g* for 2 minutes and resuspended in a microfuge tube in 1mL extracellular solution (described below) with 10 mM D-glucose and rotated at RT until use.

### Electrophysiology

Whole-cell voltage-clamp recordings were performed at RT (21-22°C) in an RC-24N recording chamber fixed to a glass coverslip (Warner Instruments), mounted on a Zeiss Axiovert 35 microscope illuminated with a Zeiss HBO 100W AttoArc lamp and filter set for epifluorescent detection of GFP expressing cells. Approximately 40 μL of cell suspension was added to the pre-filled chamber and allowed to adhere to the glass bottomed chamber for 2-10 minutes. Fresh external solution was perfused through the chamber prior to patching. Borosilicate pipettes (1.5 mm OD, 0.86 mm ID, Sutter instruments cat # BF150-86-7.5HP) were pulled, fire-polished, coated with Sylgard. Tip resistances were 1-2 MΩ, when filled with the internal recording solution. GFP expressing cells were patched and signals were amplified with an Axon Axopatch 200-B (Molecular Devices) and acquired with an Instrutech LIH 8+8 ADC board (HEKA). GΩ seals were obtained, and pipette capacitance was corrected for prior to break-in achieved by suction. Access resistance (R_s_) was typically 1-4 MΩ. 60%-80% R_s_ compensation (10 μs lag) and prediction was used to reduce voltage error to less than 10 mV as determined from the product of the peak current and R_s_ with compensation. P/5 leak subtraction protocol was used during recording. Signals were pre-filtered with a low-pass Bessel filter at 5 or 10 kHz before digitizing at 20 kHz and recorded with Patchmaster (HEKA, version 2×90.2) on a Windows 7 PC. The solutions were as follows in mM: External 3.5 KCl, 155 NaCl, 10 HEPES, 1 MgCl_2_, 1.5 CaCl_2_ adjusted to pH 7.4 with NaOH, and 315 mOsm; Internal: 35 NaCl, 70 CsCl, 50 CsF, 1 EGTA, 10 HEPES adjusted to pH 7.4 with CsOH at 310 mOsm. After break-in, cells were held at −120 mV and tested for stable Na^+^ current with depolarizing 35 ms voltage steps to −10 mV from −120 mV collected every 5 s for up to 5 minutes to allow for a stable level of current prior to vehicle addition. Once stable current levels were achieved, 150 μL of vehicle was manually added to the bath with displaced solution removed via an overflow vacuum line. After approximately 5 minutes, whole cell parameters were checked, and toxin (described below) was added by the same method as vehicle. Once apparent block plateaued, whole cell parameters were checked and adjusted as necessary, and pulsing resumed. To measure dissociation, gravity-fed perfusion with fresh external solution was started at a rate 1-2 mL/min during recording. Cells with stable leak and R_s_ allowing fitting to a single-exponential function (see below) throughout the experiment were included for analysis.

### Toxin preparation

Lyophilized WT-KIIIA was purchased (Alomone labs, Jerusalem, IS), reconstituted in water and stored as 100 μM stock aliquots at −80°C prior to use. Toxin variants were produced by solid state synthesis as described previously (Zhang et al., 2007) and stored as stock aliquots at −80°C prior to use. Stock concentrations were checked by 280nm absorbance on a Nanodrop 2000 spectrophotometer (ThermoFisher) with extinction coefficients determined by the ExPASy ProtParam online tool (Gasteiger et al., 2005). Stock aliquots of toxin were diluted in equal volumes of 2x External solution with 0.2% BSA for working solutions of toxin in vehicle of 1x External solution with 0.1% BSA and further diluted in 1x vehicle to the working concentration. Vehicle for controls were prepared in the same manner.

### Modeling and simulation analysis

Structural modeling data were analyzed using Rosetta and rendered using UCSF Chimera (Pettersen et al., 2004), VMD (Humphrey et al., 1996) was used to analyze MD simulation data. All data were plotted in R using ggplot2 (Hadley, 2016).

#### Tunnel detection for KIIIA block (*Figure 1C*)

We used CAVER (version 3.0) (Chovancova et al., 2012) to detect tunnels passing by KIIIA. Coordinates of Lys 7 in KIIIA were used as a searching starting point with probe_radius 0.9, shell_radius 5.0, shell_depth 4.0 and max_distance 10. Multiple tunnels were detected for the whole structures. We visually select only tunnels that have maximum radii greater than 2 and neighboring KIIIA for presentation.

#### Fractional contacts (*Figure 5*)

Fractional contact is defined as probability of finding two residues, one on the KIIIA and one on the channel forming contacts over time course of simulation. We considered two residues are in contact if any heavy atoms of one residue is within 4 Å of any heavy atoms of the other residues. Only contacts that have probability greater than 0.25 are shown for clarity.

#### Interface RMSD (*Figure 5 – figure supplement 1*)

We used 10 Å as a cutoff for interface calculation between KIIIA and the channels. The interface is comprised of the KIIIA itself and channel residues that are within 10 Å of KIIIA heavy atoms, defined at the beginning of the simulations. Backbone heavy atoms of the interface were used for RMSD calculation.

### Electrophysiology analysis

Electrophysiology data were analyzed and plotted in IGOR 7 Pro (Wavemetrics). Geometric means of kinetic parameters were determined using Excel (Microsoft) and plotted in IGOR 7 Pro. Curve fitting was performed in IGOR Pro 7 as described previously (Dockendorff et al., 2018). To determine time constants of toxin association and dissociation (*τ_on_* and *τ_off_*, respectively), peak currents during depolarizing voltage steps were plotted by time, and data were fit with a single exponential function:

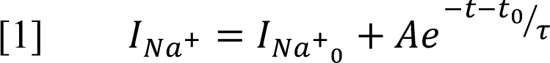

The association rate *k_on_* was determined by equation 2, or equation 3 (McArthur et al., 2011) was used for KIIIA-variant H12A where the maximal block at saturating concentrations (*F_block_*) was already known and *k_off_* could not be determined independently:

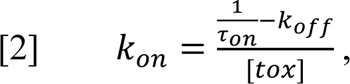

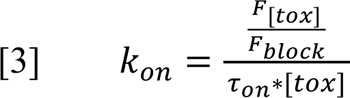

*k_off_* was determined by equation 4:

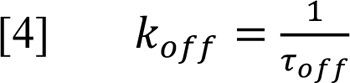

Affinity was determined kinetically as the dissociation constant *K_d_* via equation 5:

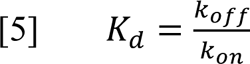

The slow dissociation of WT-KIIIA from WT-hNa_V_1.7 made measurement of *k_off_* difficult due to limited recovery during the experiment, thus values shown here are estimates assuming eventual full recovery of the maximal current before toxin association. The resulting affinity values are consistent with previous reporting of kinetic determination of affinity for this channel (McArthur et al., 2011). Maximal block and IC_50_ for WT-KIIIA x WT-hNa_V_1.7 was determined from concentration-response of peak residual current at equilibrium with equation 6. The Hill coefficient *h* was assumed to be 1 in accordance with a single binding site:

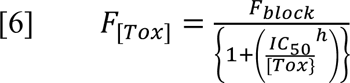

Maximum fractional block at saturating concentrations was determined from kinetic data and observed block (*F_[tox]_*) for other channel and toxin variants, except where noted, according to equation 7 (McArthur et al., 2011):

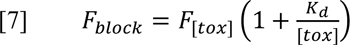

The low affinity of H12A hindered precise measurement of dissociation kinetics; both the rapid rate of dissociation and the low degree of block limited the number of data points for fitting a single exponential to the dissociation data. The dissociation rate is extrapolated from Fractional block assuming maximal block at saturating concentration of 0.877, as reported previously (McArthur et al., 2011). The E919Q x D11A condition was affected by the same difficulty, thus kinetic values reported assumed similar levels of block to those observed during the E919Q x WT-KIIIA condition (0.92, Table 3). The lack of effect of KIIIA-D11A on channel block suggests that any additive effect of the E919Q x D11A double mutant condition would not reduce the level of block seen from WT-KIIIA on Na_V_1.7-E919Q. Any errors in estimation of maximal block with toxin variants or channel mutants did not affect the calculation *K_d_* = *k_off_* / *k_on_*, and thus did not affect the coupling coefficients and energies calculated for double-mutant cycle analysis which are derived from *K_d_* = *k_off_* / *k_on_*.

Coupling coefficients (Ω) were calculated from the dissociation constants of the four conditions for each cycle according to equation 8 (Hidalgo and MacKinnon, 1995), where “*K_d_wm*” would represent the dissociation constant for WT-hNa_V_1.7 x Toxin-variant condition:

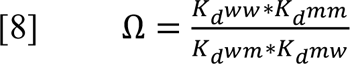

### Descriptive statistics

Arithmetic means and standard error were calculated for *F_block_*, while logarithmically scaled kinetic parameters were summarized with geometric means and standard deviations. Standard errors of kinetic parameters were obtained for the tables as the dividend of the standard deviation and the square root of the sample size for each toxin-channel pair, as noted in parentheses in each table. Errors for coupling coefficients and coupling energies were calculated by linear propagation of error from fractional standard deviations of the reported *K_d_* values for the toxin – channel mutant pairs used to calculate the coupling coefficients.

---

**Figure 1—figure supplement 1.**
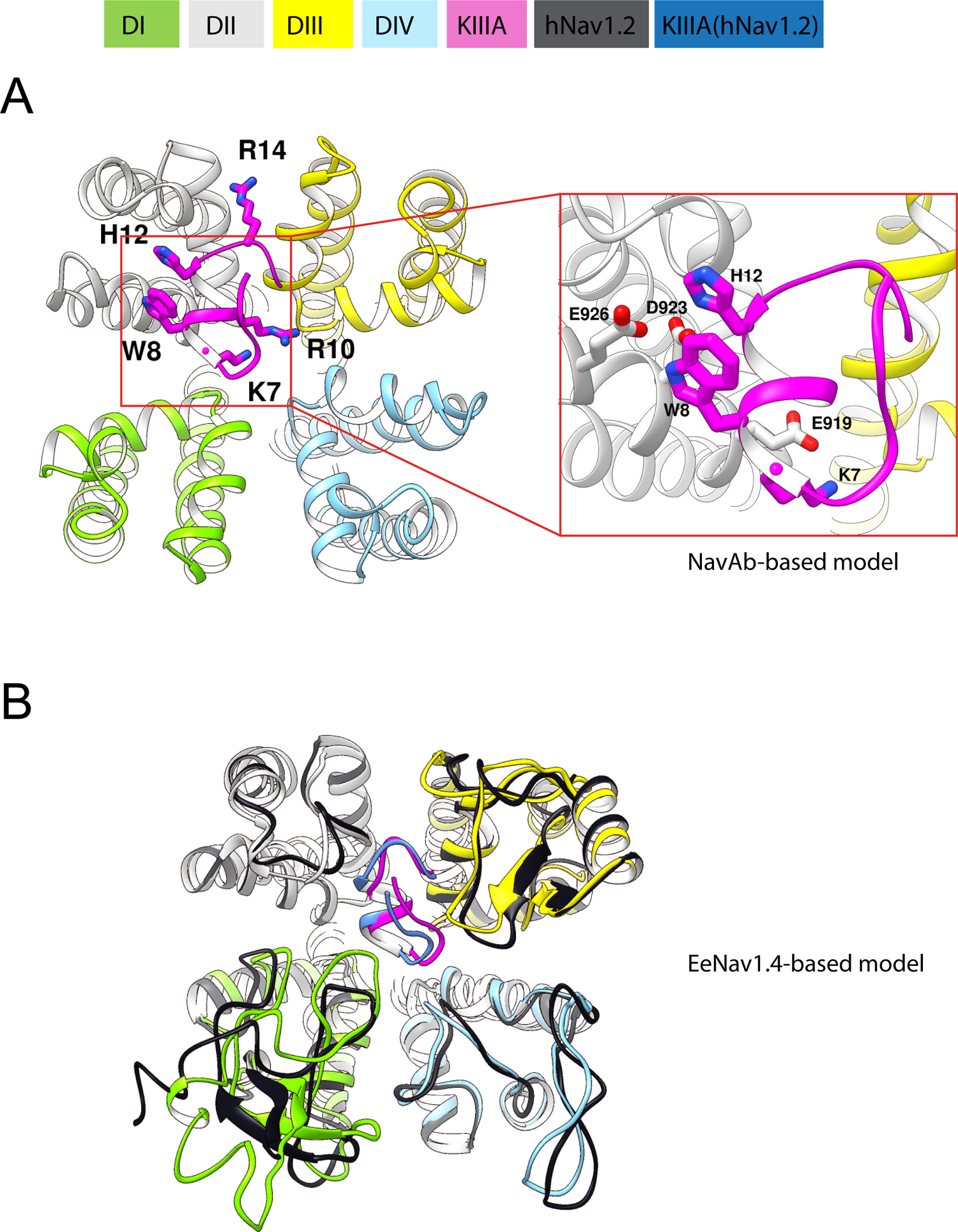
Rosetta models of hNa_V_1.7 – KIIIA complex. Domains are depicted according to color keys. (**A**) Extracellular view of the pore region of the Rosetta model of the KIIIA - hNa_V_1.7 complex based on bacterial Na_V_Ab structure (Payandeh et al., 2011). (**B**) Extracellular view of the pore region of the KIIIA - hNa_V_1.7 model based on eeNa_V_1.4 structure (Yan et al., 2017) in superposition with the cryo-EM structure of the KIIIA - hNa_V_1.2 complex (Pan et al., 2019).

**Figure 2—figure supplement 1.**
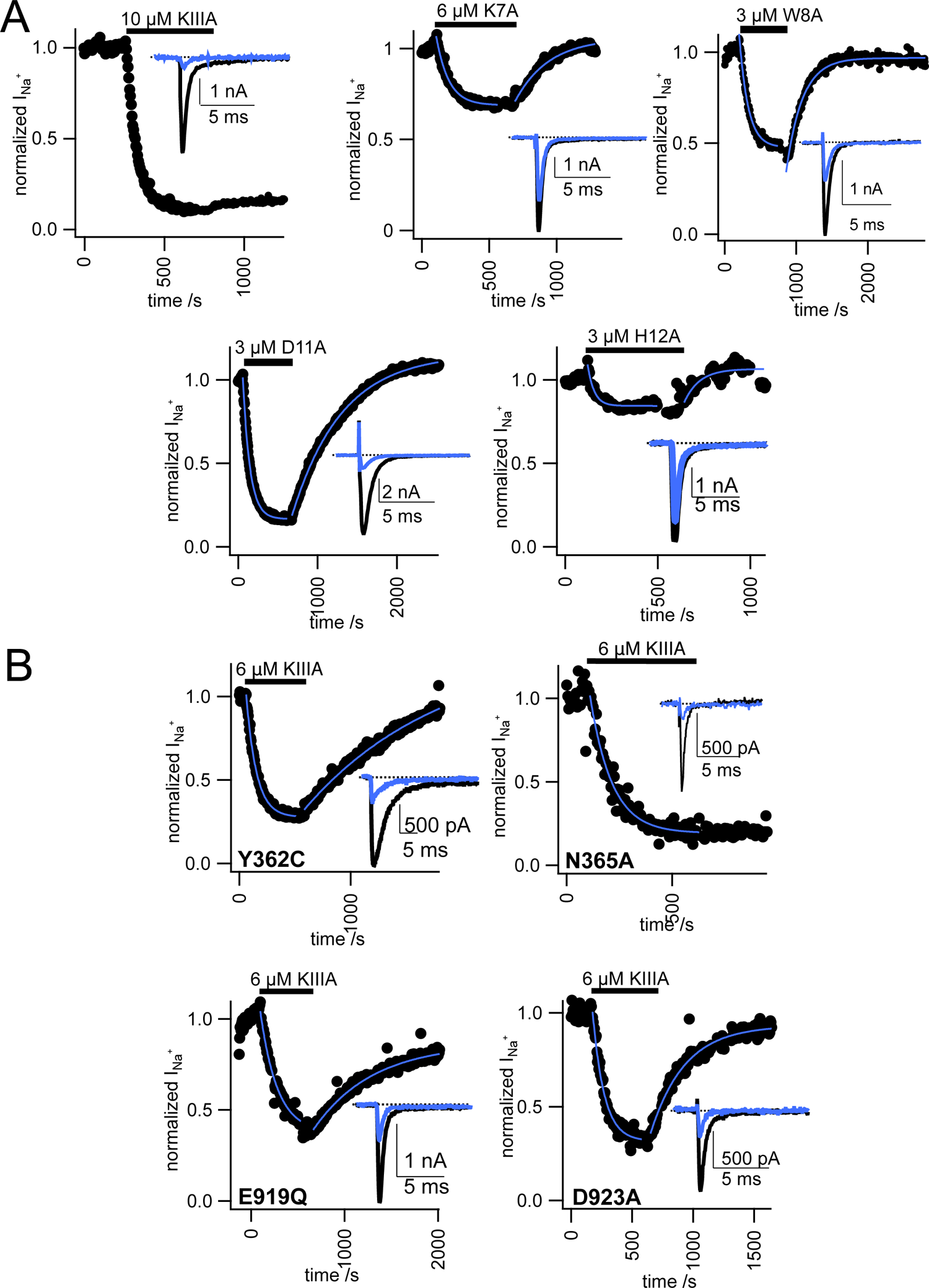
Representative data from functional studies of toxin variants and channel mutations. (**A**) Normalized peak I_Na_+ from whole cell voltage clamp experiments with WT-hNa_V_1.7 in the presence of toxin variants as stated in each plot, exponential fits of association and dissociation shown in blue. Raw current traces (inset) before toxin (black) and after toxin addition (blue). (**B**) Normalized peak I_Na_+ from whole cell voltage clamp experiments with hNa_V_1.7 mutants as labeled in the presence of WT-KIIIA as in panel A.

**Figure 3—figure supplement 1.**
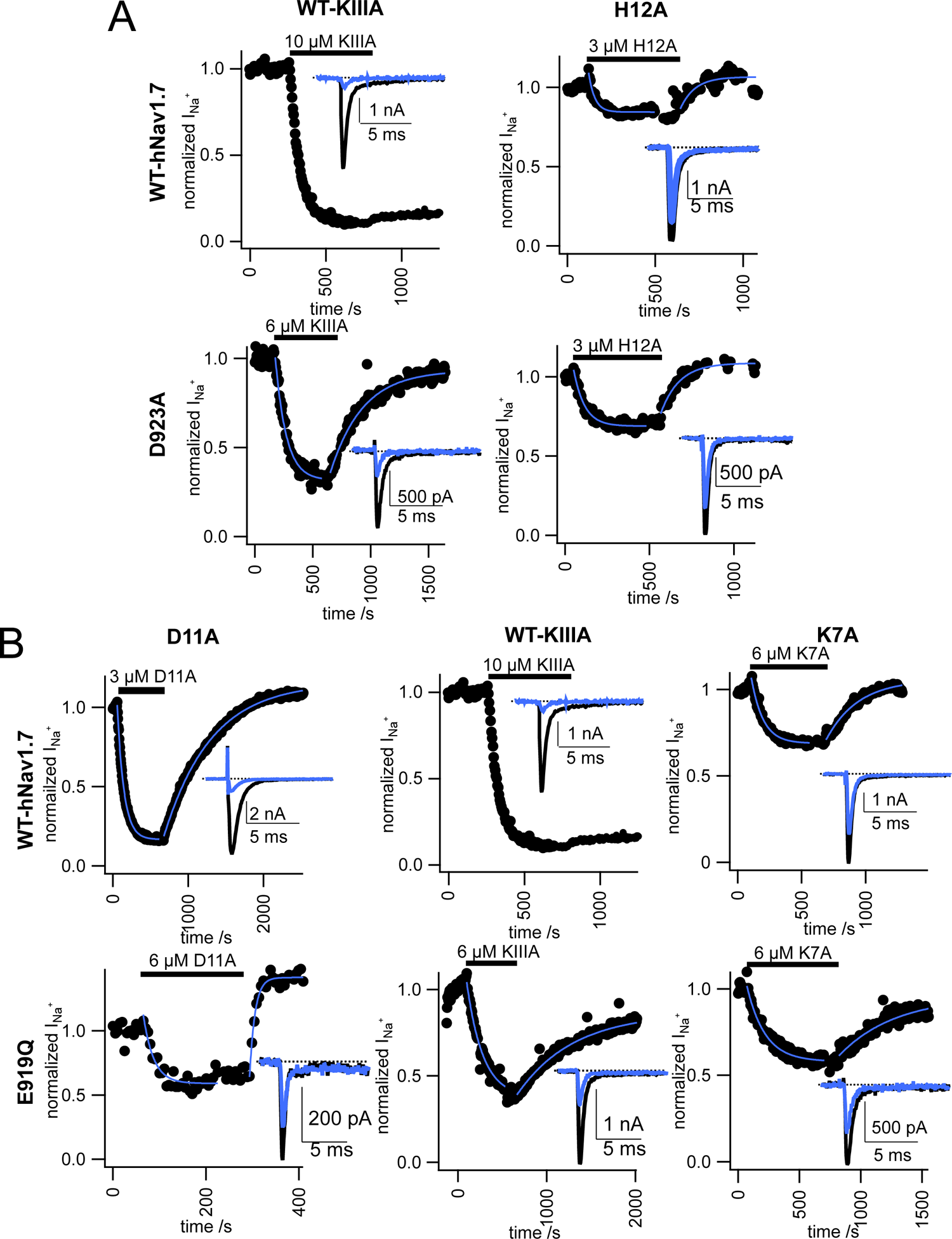
Representative data from double-mutant cycle analysis. (**A**) Normalized peak I_Na_+ from whole cell voltage clamp experiments with WT x WT, single, and double mutations D923A and H12A as indicated by column and row labels. Exponential fits of association and dissociation shown in blue. Raw current traces (inset) before toxin (black) and after toxin addition (blue). (**B**) Normalized peak I_Na_+ from whole cell voltage clamp experiments, as in panel A for the two residue pairs E919 x K7 and E919 x D11.

**Figure 4—figure supplement 1.**
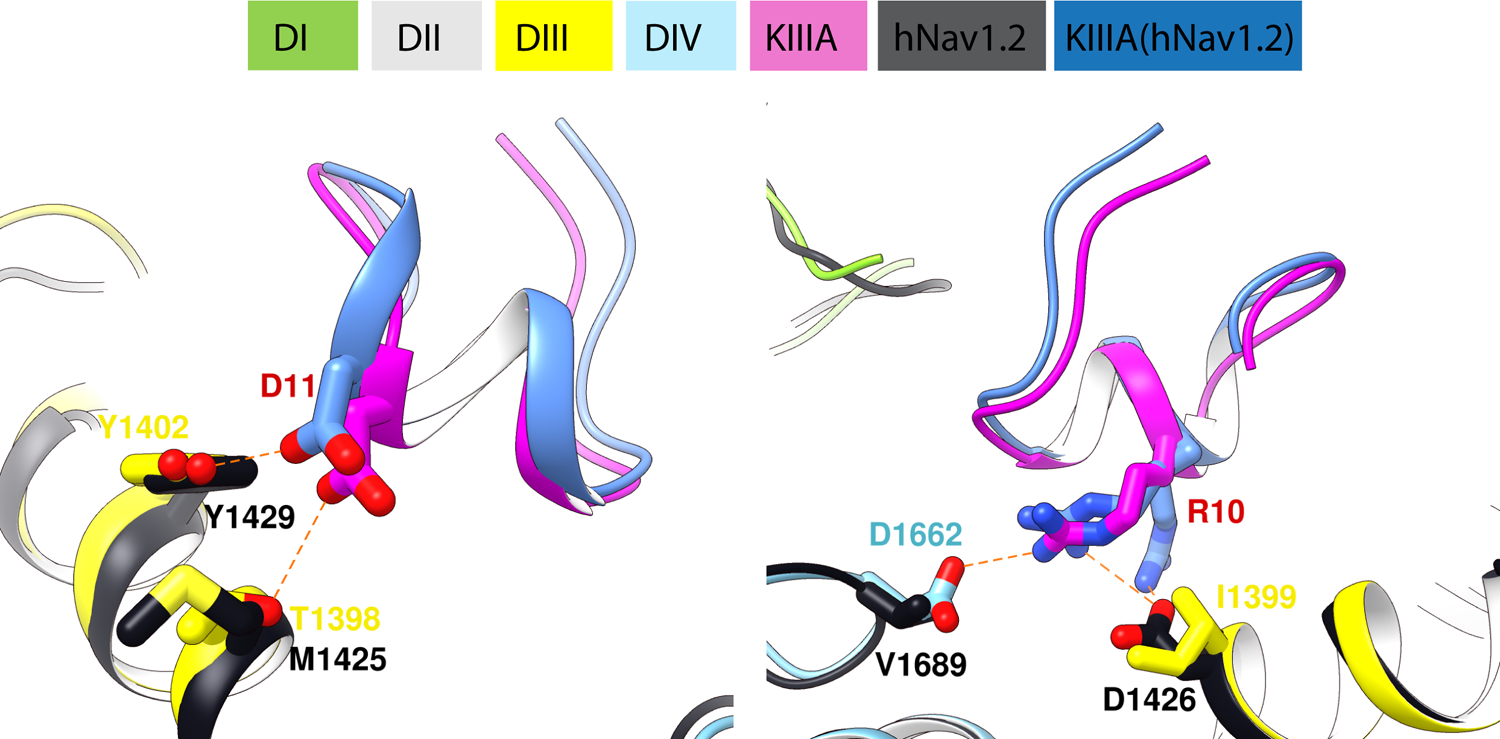
Differences in interactions of KIIIA residues D11 (left panel) and R10 (right panel) between the KIIIA - hNa_V_1.7 model and the KIIIA - hNa_V_1.2 structure. Both alternative sidechain rotamers of KIIIA R10 residue identified in the KIIIA - hNaV1.2 structure are shown. Domains and labels are depicted according to color keys.

**Figure 5—figure supplement 1.**
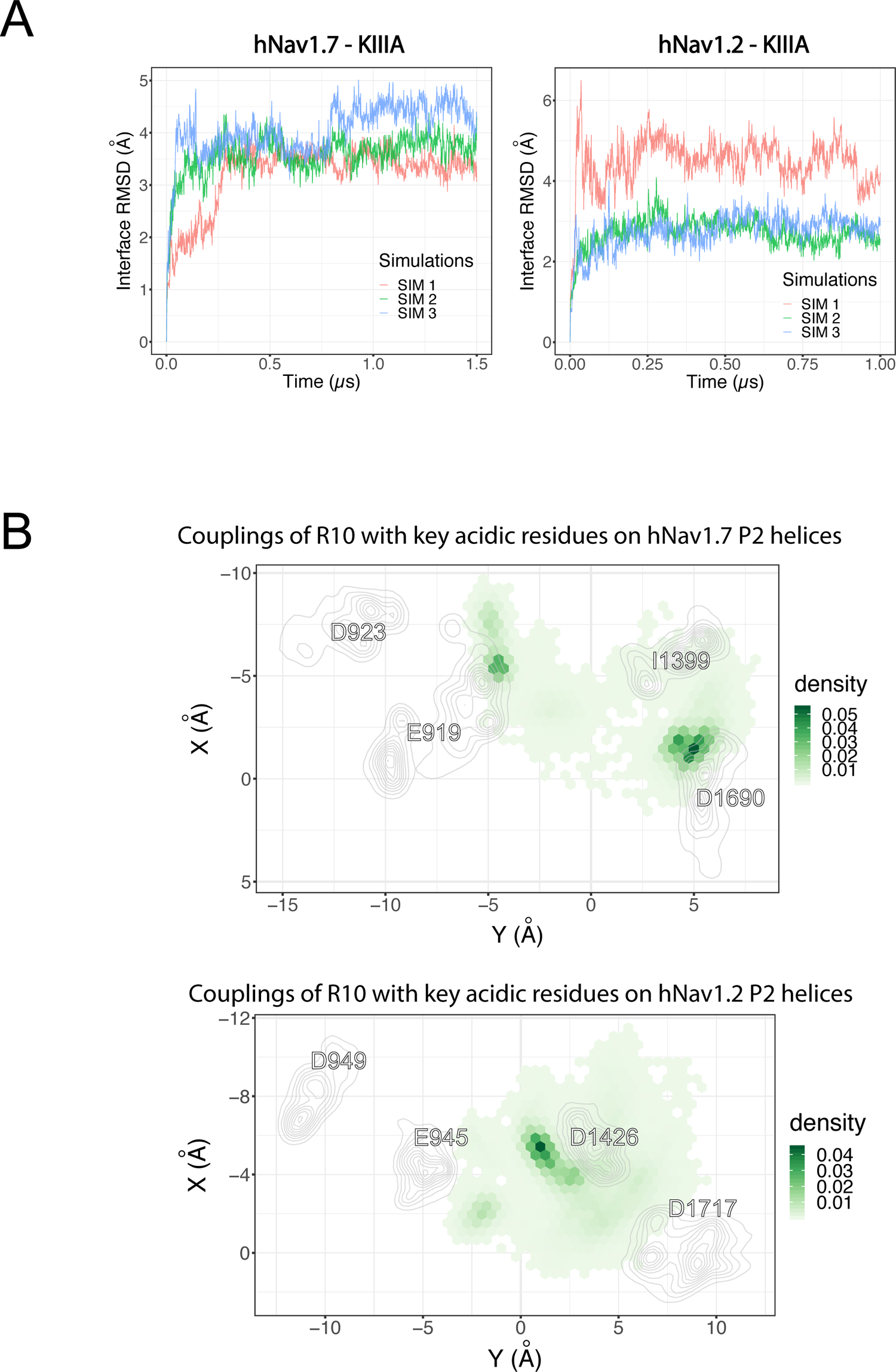
MD simulation of KIIIA - hNa_V_1.7 model and KIIIA – hNa_V_1.2 structure. (**A**) Interfacial RMSD of the KIIIA – hNa_V_1.7 and KIIIA – hNa_V_1.2 complexes in different independent simulations. (**B**) Couplings of R10 with key acidic residues on the P2 helices of hNa_V_1.7 (upper panel) and hNa_V_1.2 (lower panel) are depicted as density projections of R10 and the acidic residue on P2-helix (labeled) on the XY plane. For clarity, densities of the acidic residues are shown as contour plots using kernel density estimation.

**Figure 5—figure supplement 2.**
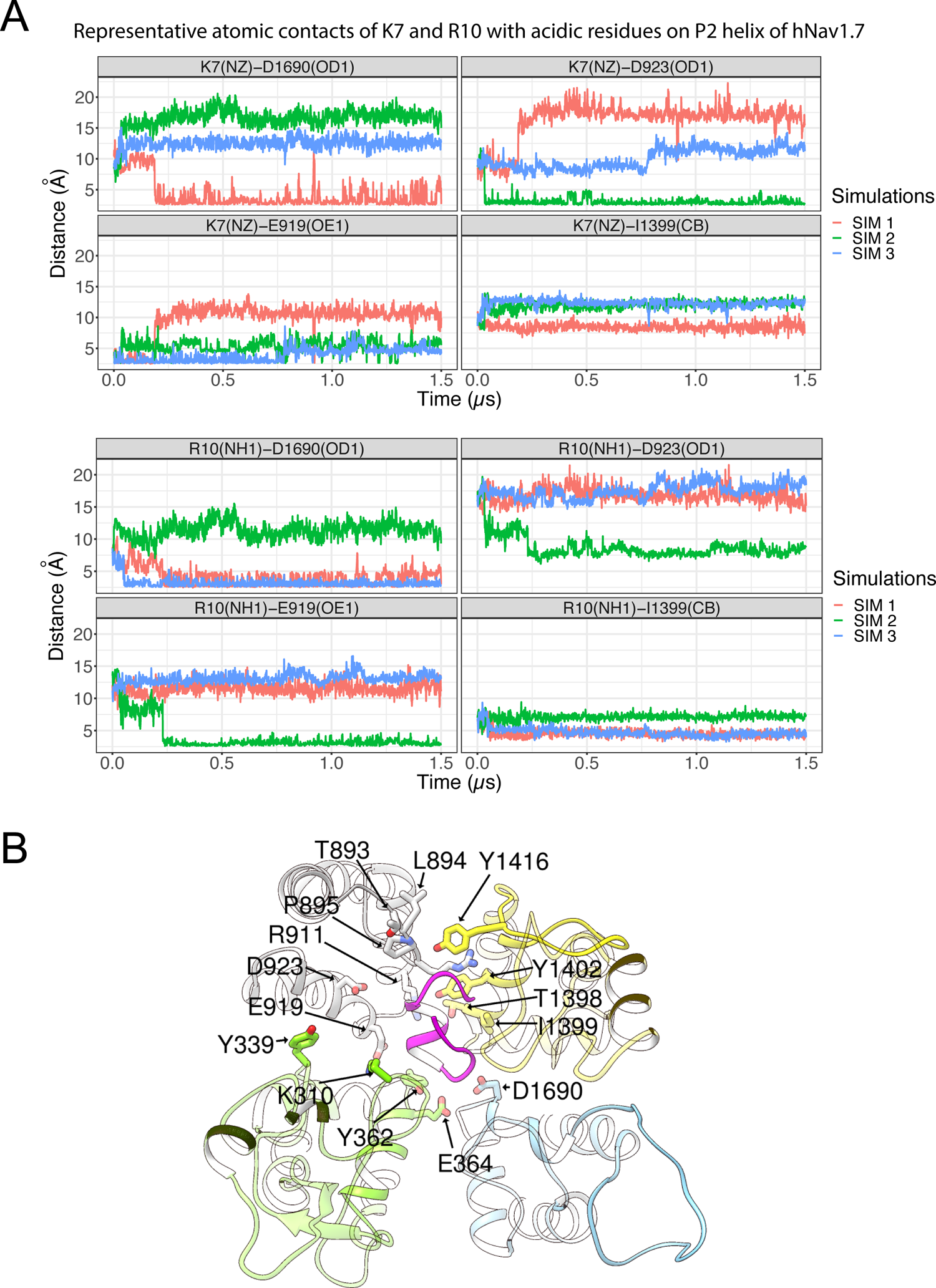
MD simulation of KIIIA - hNa_V_1.7 model. (**A**) Time series of representative contacts of K7 and R10 with acidic residues on P2 helix of hNa_V_1.7. Labels show residue numbers and their associated atom names (in parenthesis). (**B**) A snapshot of the last frame of the 1μs simulation. Domains are depicted according to color keys and channel residues that have high fractional contacts in Figure 5A are shown in stick representation and labeled.

**Figure 5—figure supplement 3.**
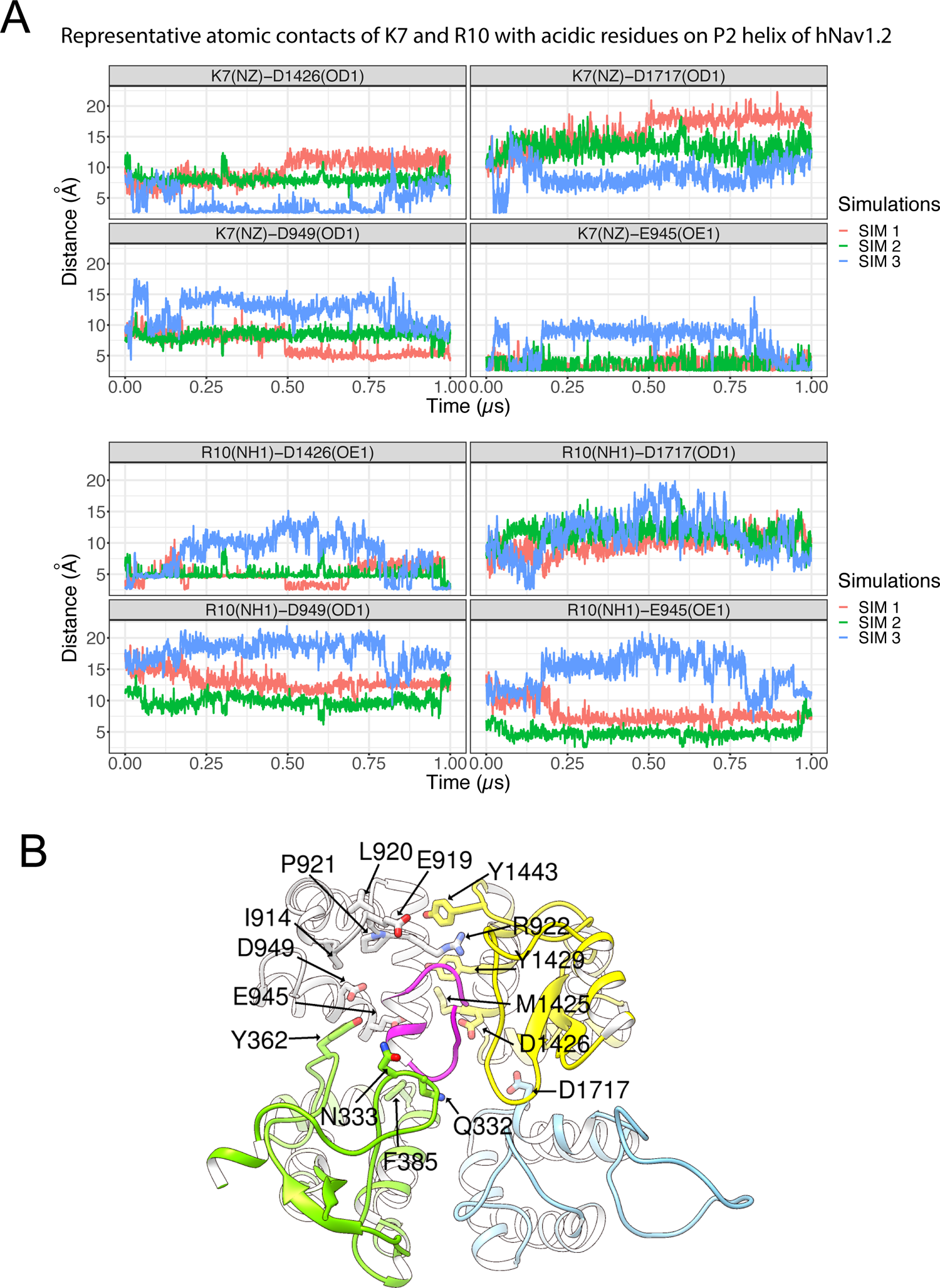
MD simulation of KIIIA - hNa_V_1.2 model. (**A**) Time series of representative contacts of K7 and R10 with acidic residues on P2 helix of hNa_V_1.7. Labels show residue numbers and their associated atom names (in parenthesis). (**B**) A snapshot of the last frame of the 1μs simulation. Domains are depicted according to color keys and channel residues that have high fractional contacts in Figure 5B are shown in stick representation and labeled.

